# Neural Geometric Representations of Social Memory for Multi-individuals in the Medial Prefrontal Cortex

**DOI:** 10.64898/2025.12.13.691981

**Authors:** Tianyu Li, Xingzhi He, Xiang Gu, Xiaohui Zhang

## Abstract

Social memory, an ability to recognize acquainted individuals and related experience, is mainly stored in the medial prefrontal cortex (mPFC). However, it remains unknown how the mPFC encodes the memory of interacted multi-individuals. We, here, report that tuned low-dimensional, geometric subspaces in population neural activity of mPFC neurons encode this form of social memory. Different social experience, in which mice encounter four conspecifics associated with reward, aversive or neutral stimulus, respectively, substantially enhances segregation among identity-specific neural geometric subspaces. Progressive recruitments of mPFC neurons conjunctively encoding social identity and associated social valences underlie the enhanced neural representations of different individuals. Thus, our findings elucidate that tuned neural subspaces in mPFC population activities stably encode and maintain the memory of acquired multi-individuals over experience.

## Main Text

Social recognition, an evolutionarily conserved brain function, is fundamental to survival and reproduction across species. Studies using non-invasive techniques have demonstrated that human and other primates strongly rely on conserved sensory cues (particularly visual) and integrated neural networks to recognize conspecifics(*1-4*). Rodents are also social animals, possessing many social behaviors such as communal nesting and cooperative care(*5, 6*).

Conventional research on social memory in mice has largely relied on the three-chamber assay, designed to assess binary discrimination between familiar and unfamiliar individuals or different emotional states(*7-13*). However, this assay often conflates novelty-driven exploration with genuine social recognition, and thus has failed to disentangle neural representations of social identity from confounding factors such as spatial location(*8, 9, 14, 15*). Furthermore, this two-choice design also has lacked of adequately reproducing social environments in nature, where individuals ought to distinguish among multiple conspecifics. Thus, whether mice could discriminate among three or more individuals and relative mechanism is implemented in the brain remain poorly understood.

Several key brain regions, including the hippocampus(*10, 13, 16-21*), nucleus accumbens (NAc)(*8, 21*) and medial prefrontal cortex (mPFC)(*6, 8, 9, 21-28*), are known to underlie the formation or maintenance of social memory. For example, the mPFC has been shown to exert multiplex functions in processing social information, such as dynamically encoding social interactions(*9*), storing social experience(*28*) and gating normal social behaviour(*25*). Yet, spiking activity responses of individual mPFC neurons, recorded from the mice in those binary social tasks, were tuned to mixed social conspecifics, positions or other social-related cues(*14, 15, 23, 29, 30*), and these results can be hardly used to resolve the neural activity mechanism for encoding distinct social individuals. While several studies have explored multi-mouse social interaction and recognition(*16, 24*), how mPFC neuronal activities encode individual social identities and the activity representations evolve with social experience remain unclear.

In the present study, we developed a novel one-*versus*-four social discrimination task that extends behavioral and neural analyses beyond the conventional one-*versus*-two (binary) comparison. Using chronic *in vivo* electrophysiological recordings with multi-wire arrays in free-moving mice, we identified distinct classes of mPFC neurons exhibiting tuned spiking response to distinct social demonstrators, spatial locations, or conjunctive social-spatial features. We further developed a cross-session alignment approach using the regularized linear regression(*31, 32*) and joint dimensionality reduction(*33*), revealing these stable, low-dimensional neural subspaces that uniquely encode individual demonstrators. These subspaces underwent experience-dependent reorganization and underlay shifts of social preference to demonstrators after conditioned social experience in the test mice. Thus, our findings reveal a mechanistic basis for the function of social discrimination among multiple social individuals.

### mPFC Neurons Exhibit Diversely Selective Responses in the Multi-Individual Social Test

The widely-used three-chamber test paradigms have generally failed to account for the modulation of neuronal activity by spatial variables and mixed selectivity (*8, 9, 14, 15, 29, 30*), despite the fact that a substantial proportion of mPFC neurons encode a combination of social and location information(*8*). To exclude spatial influence and determine whether mice can discriminate among multi-social demonstrators at the single-neuron level, we performed *in vivo* electrophysiological recordings in the mPFC using a 32-channel micro-wire array (Fig. 1A-C and fig. S1) in the newly-designed one-*versus*-four social exploration task, where a male mouse freely investigated to four male stranger demonstrators resided at four corners in a square open-field arena, respectively, for 10 min. In each day the same test mice underwent two consecutive test sessions, between which positions of paired demonstrators were switched (Fig. 1A and movie S1 to S2) to dissociate the social and spatial location information of the demonstrator. In the task, the test mice spent around 60% of the total time dwelt in the social investigation zone (Fig. 1B), in which about 53% was associated with specific social investigation behaviors such as sniffing individual demonstrators. In this study, we recorded a total of 1800∼ single neuronal units from 8 test mice, and only those mice yielding > 20 single-units per day (70% of recorded mice) were included for further analysis (Fig. 1C).

**Fig. 1.**
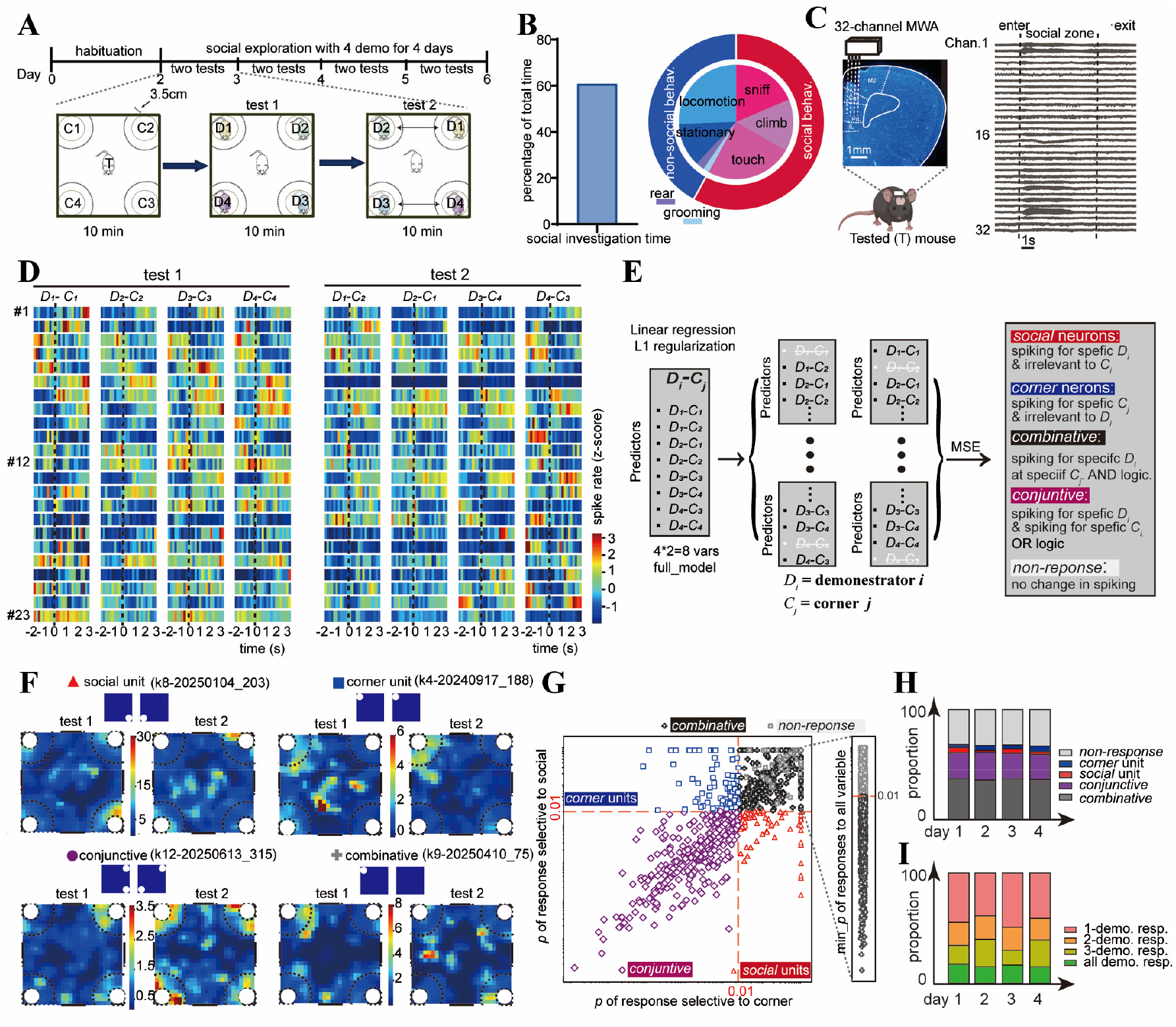
Neural response features of individual mPFC neurons in the multi-social test. (**A**) Schematic of the four-demonstrator social exploration test. (**B**) The proportions of a test animal’s time spent in the social zone (left) and related social exploration behaviors (right, red) as well non-social explorations (blue) in the 10 min test time. (**C**) *Left*: Schematic illustration of implantation of a 32-channel micro-wire array (MWA) in the mPFC. *Right:* Raw traces of *in vivo* recorded neural activities from individual 32 channels, covering a complete time duration spent in the social zone (dotted line: time points for entering and exiting, respectively). (**D**) Changes of spiking rates (*z*-scores) of representative mPFC single-units in the social zone in the consecutive social test 1 and 2 sessions, respectively. Dotted grey lines indicate the time of entering the social zone. (**E**) A diagram illustrating the procedure of utilizing the Lasso Regression to classify neuronal units encoding distinct spatial or social features. D_*i*_and C_*j*_depict the demonstrator mice *i* and the corner *j*, respectively. (**F**) Representative *social* or *corner* neuron, *conjunctive* neuron and *combinative* neuron, respectively from the analysis in the **e**, indicated by distinct heat-maps of their mean spiking rates at different arena locations between the social test 1 and 2 sessions. (**G**) Distributions of *p*-values for clarifying the featured *social* neurons (red triangles), *corner* neurons (blue squares), *conjunctive* neurons (purple circles), *combinative* neurons (dark grey), and *non-response* units (light grey), based on the criterion of *p* < 0.01. (**H**) Changes of the proportions of distinct featured units across the 4 days of social test. (**I**) Changes of the proportions of social units encoding demonstrator 1, 2, 3 and 4, respectively, across 4 test days.

Based on differential changes of spiking rate of recorded units during animal dwelling within the social zone in the two session tests (Fig. 1C), we adopted the regularized Lasso Regression model using social identity (*D*_*i*_) at its corner position (C_*j*_) as predictors(*31, 32, 34*) to determine the unit’s responses to either social or spatial features (Fig. 1E). Significance and generalizability of the model were assessed via a nested 10-fold cross-validation, testing whether the removal of a given predictor caused a significant increase in the mean squared error (MSE)(*15*). With the latter approach, we could categorize the units into five groups based on their distinct response selectivity: (1) *social* neurons: spiking rates were selectively modulated by specific *D*_*i*_ in both test sessions (Fig. 1F upper left and Fig.1G red; 3.2% of total units, shuffle = 0%); (2) *corner* neurons: rates selectively modulated by specific *C*_*j*_ in both sessions (Fig. 1F upper right and Fig. 1G blue; 4.5%, shuffle = 0%); (3) *Conjunctive* neurons: changes were tuned to either *D*_*i*_ or *C*_*j*_ across sessions (Fig. 1F bottom left and Fig.1G purple; 24.4%, shuffle = 0%); (4) *Combinative* neurons: rate changes exclusively to a specific *D*_*i*_ in a specific *C*_*j*_ (Fig. 1F bottom right and Fig. 1G black; 35.7%, shuffle =1.3%); and (5) *non-response* neurons: no tuned rate response to any variable in the social zone (32.3%, shuffle =98.7%). Consistent with the previous findings(*8, 9, 14, 15*), these results indicate that many mPFC neurons exhibit their spiking selectivity to multiple cues and a large portion of them act as the *combinative* or *conjunctive* encoding neurons(*8*). We note that the overall proportions of individual functional neuron subtypes, recorded by all test mice, did not vary much across the 4 days of social exploration (Fig. 1H).

Among 521 neurons that encoded social individuals (including the *social* neurons and *conjunctive* neurons), we further quantified their spiking specificity to either single or multiple demonstrator mice. The results showed that about 44.1% of them exhibited their selectivity exclusively to single demonstrator, while the remaining 55.9% encoded 2-4 demonstrators (Fig. 1I). The proportions of social neurons encoding single or multiple individuals showed not significant changes across the 4 days.

Taken together, our recording results suggest that the majority of responsive neurons exhibit multi-selectivity or conjunctive tuning, while a small proportion of mPFC neurons specifically encode single or multiple social identities of four interacted individuals. These findings underscore the notion that mPFC neurons integrate diverse sensory inputs and behavioral variables, and their spiking responses display mixed-tuning properties(*8, 14*). Moreover, the mixed identity tunings of recorded mPFC social neurons implicate that social identity may be encoded and maintained in a distributed pattern, an efficient strategy that allow to maximize the capacity of storing complex social information among mPFC neurons(*14, 30*).

### Population Activity Coding of Multiple Social Identities

Because the mPFC functions as an integrative and hub-like region for processing various information in social recognition(*2, 6, 8, 9, 12, 22, 23, 25, 28, 35*), we next examined whether population activity of mPFC neurons underlies social discrimination among four demonstrators. To decode which social demonstrator the test mice were encountering, we trained Support Vector Machines (SVMs) with 80% of the recorded unit activity patterns and evaluated their performance on the held-out 20% set with either a linear or non-linear radial basis function (rbf) kernel (Fig. 2A). The performance accuracies of both linear and non-linear SVM classifiers were significantly above the chance level (F1_score_linear_= 38.6% ± 6%, F1_score_rbf_= 45.1% ± 9%, shuffled accuracy: 25.0% ± 2.5%), and higher F1-scores were found in the rbf kernel (Fig. 2B to C). These results indicate that population activity in the mPFC reliably encodes social identity and such social representations are inherently nonlinear. The prevalence of *combinative* and *conjunctive* neurons in the mPFC could partially account for the observed nonlinearity in neuronal representation. Moreover, F1-scores of the rbf kernel SVM varied little in discriminating individual conspecifics across multiple 4 days of recording (Fig. 2D), suggesting a robust presentation by mPFC population neuronal activity.

**Fig. 2.**
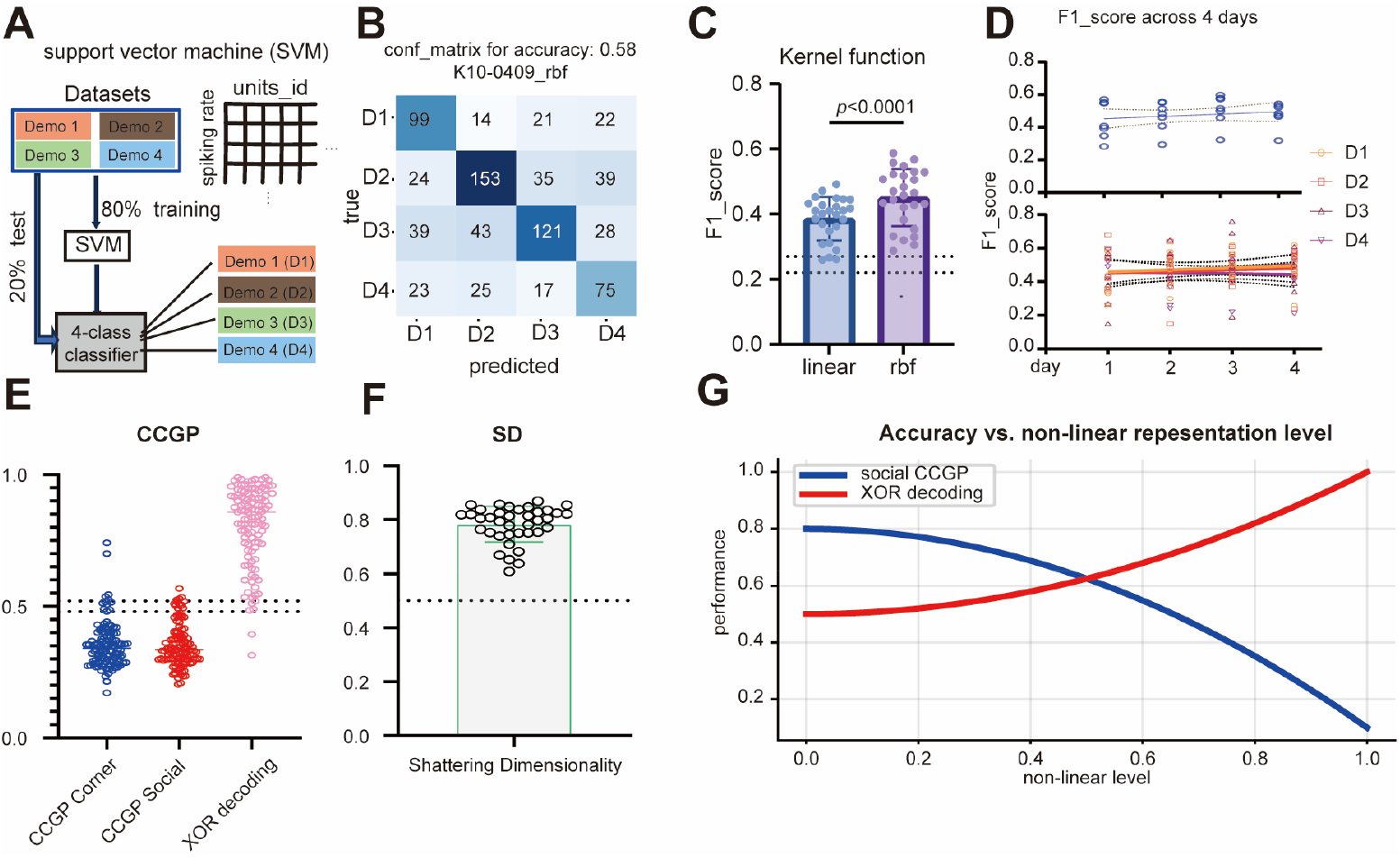
Decoding multiple individuals based on mPFC population neuronal activities. (**A**) Schematic diagram illustrating the use of support vector machine (SVM) classifier to test the accuracy in discriminating multi-individuals (Demo *i*) on the basis of neuronal spiking activities recorded during the test-mouse dwelling within the social zone. Spiking trains of all neurons were binned with a 250-ms sliding window, and 80% and 20% were used for the classifier training and test, respectively, in which all class were balanced. (**B**) An example of the confusion matrix calculated from the SVM model with radial basis function (rbf) kernel. (**C**) Comparison of calculated F1 scores from the SVM classifiers with a linear or a rbf kernel (t-test). Dotted lines: the shuffled F1 score (0.25±0.04). (**D**) Changes of the averaged F1_score (upper) or the individual F1_scores (bottom) of the SVM for classifying distinct demonstrators across days. Dots represent individual tested mice. (**E**) Results of calculated Cross-Condition Generalization Performance (CCGP). Note the values for the corners and social demonstrator are substantially lower than the null model (0.50±0.02), while the trial decoding (XOR) accuracy are higher than the null model (0.50±0.02). (**F**) the Shattered Dimension (SD) value is greater than the null model (0.50±0.02). (**G**) the Hopfield Recurrent Neural Network simulations showing the opposite changes of the CCGP of social identity decoding (blue) and XOR decoding (red) when the nonlinearity of neural representations gradually increases.

To assess the nonlinearity of social representations in the mouse mPFC, we computed the cross-condition generalization performance (CCGP) and the shattered dimensionality (SD) of neural representation, based on the approach developed by Bernardi et al.(*36*) and Boyle et al.(*37*) (Fig. 2E to F and fig. S2, also see the materials and methods for these metric calculation). The CCGP measures the abstractness of a variable’s neural representation, defined by the ability of a linear decoder that is trained on a subset of trial conditions to generalize its classification to held-out other conditions (fig. S2), and a high CCGP value (significantly > 0.5) indicates a representational geometry, such as clustering or factorized, parallel subspaces, *vice versa*. Meanwhile, the SD measures the overall expressive capacity and linear separability of the population code, and a high SD value reflects a high-dimensional geometry that allows a linear readout to support a vast repertoire of input-output mappings, indicating computational flexibility. In our experiment in which positions of 4 social demonstrator were fully switched across the 2 consecutive exploration tests and yielded 8 unique demonstrators-in-position conditions, we found that respective values of CCGP for social and corner conditions were significantly lower than the shuffled baseline (Fig. 2E), while their SD values were significantly higher than the shuffled baseline (Fig. 2F). Meanwhile, the exclusive-OR (XOR) decoding accuracy (*36-39*) remained notably high performance (Fig. 2E), indicating pronounced nonlinear and high dimension encoding of social information in the mouse mPFC. Furthermore, simulations using a Hopfield recurrent neural network (see the materials and methods) demonstrated that as the degree of nonlinearity increased, CCGP decreased along with increased XOR decoding accuracy (Fig. 2 E to G), suggesting that social representations in the mPFC are highly nonlinear geometry(*36-39*).

Since social-related neurons (including *social, conjunctive*, and *combinative* neurons) exhibited significantly higher feature importance for decoding than non-social neurons (*corner* and *non-response* neurons) (fig. S3A), we further evaluated differential contributions of social-related neurons and non-social neurons in the decoding performance by random subsampling of equal numbers of neurons from these two groups, respectively (fig. S3B top). This revealed significantly higher decoding accuracy from social-related neurons. Conversely, randomly ablating an equal number of neurons from each group caused a more pronounced accuracy drop when social-relative neurons were removed (fig. S3B bottom). Collectively, these complementary results confirm a dominant role of social-related neurons in supporting social discrimination. Although the social-related neurons encoded more information of social identity, these observed non-social neuron also complimentarily supported social memory (fig. S3C to E), underscoring the notion that social memory is facilitated by collaborative engagements of multiple neural orthogonal principal components(*40*) (fig. S3F).

### Stable Neural Activity Subspaces for Identity Representation

We next delineated how mPFC population activity robustly encodes identities of multiple social individuals across days. First, we employed the Cebra(*41*) to make a nonlinear dimensionality reduction (multi-session mode) of population activity, recorded from different days, and found similar embedded structures for representing respective social identifies across days. (Fig. 3A and fig. S4b). Second, we developed a new method to compare embedding spaces of population activity across days, with approaches combining the regularized regression and the joint dimensionality reduction in order to overcome the possible noises brought by the mix encoding of multiple behavioral variables(*29*) and the difficulty in cross-day alignment of neuron ensembles, respectively (see the materials and methods for the dimension reduction). With this, we concatenated the regression coefficient matrices across days into a unified design matrix and performed joint singular value decomposition (SVD) and aligned all sessions into a low-dimensional subspace (Fig. 3B). We found that calculated similarities between dimensions across the tested 4 days were fair high (cosine similarity > 0.85; Fig. 3C), indicating that recorded units from different days in the same animal are embedded in a nearly identical low-dimensional subspace.

**Fig. 3.**
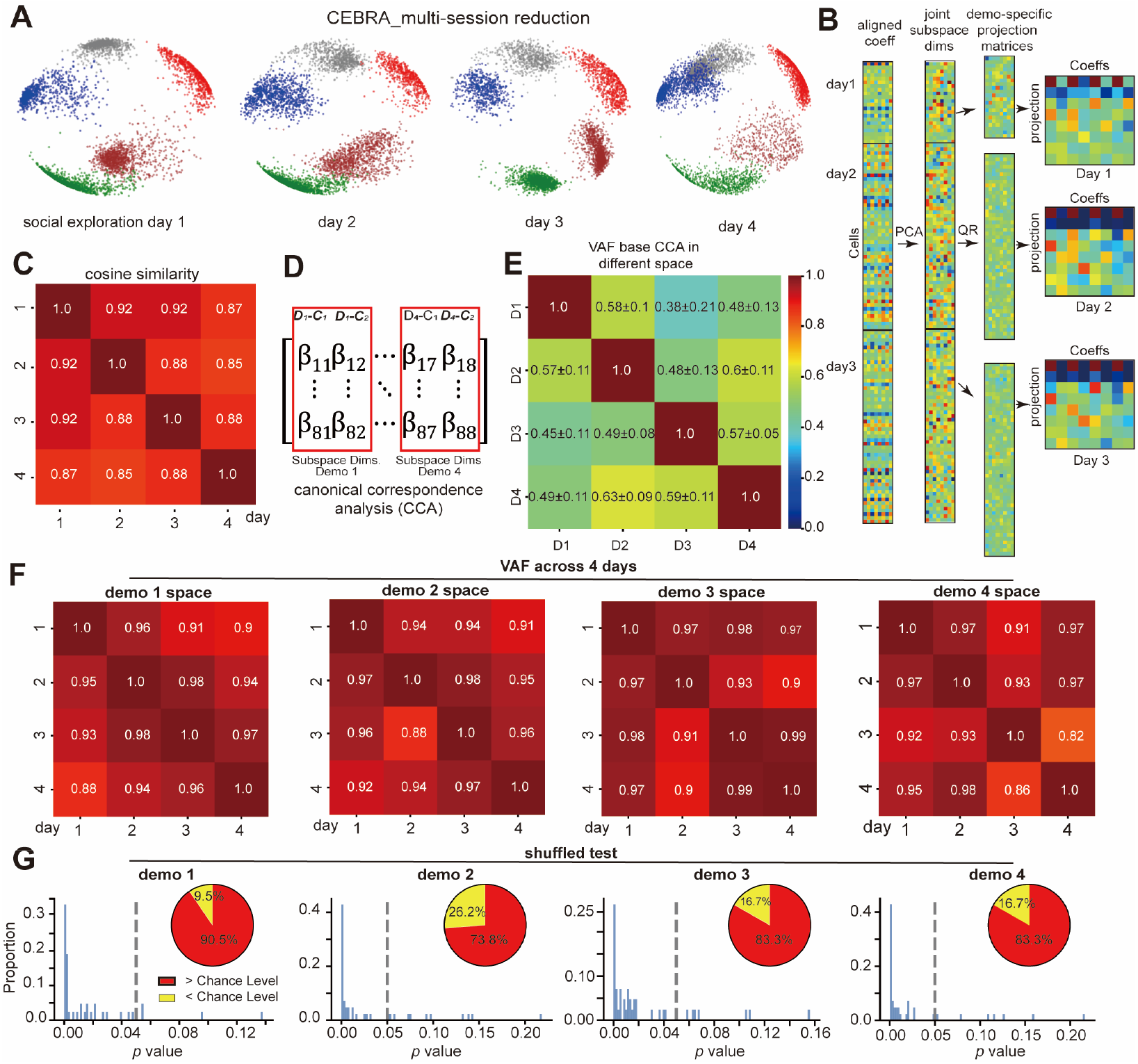
Stable neural representations of individual identity in the mPFC. (**A**) Stable cluster structures of neuronal activities in the mPFC across the 4 test days, revealed by the CEBRA multi-session reduction. Clusters in colors represent units that spiked within the social zone and encoded distinct 4 demonstrators, while clusters in grey are ones that spiked outside the social zone. (**B**) A schematic diagram illustrating a unified dimensionality reduction framework based on the joint Singular Value Decomposition (SVD). Matrices from different days were concatenated along the aligned “demonstrator × location” regression coefficient (weight) and projected into a unified low-dimensional space, in which the horizontal and vertical axis represent the coefficients and the projections as ‘super neuron’ capturing the overall encoding patterns of neuronal population. (**C**) Cosine similarity between neural representations across days, calculated from the (**B**). (**D**) A diagram illustrating the Schematic of Canonical Correlation Analysis (CCA) to analyze cross-day stability of subspaces, constructed by the dimensionality reduction of the two projection arrays, ‘D*i*-C_*1*_’ and ‘D*i*-C_*2*_’. D: demonstrator, C: corner. (**E**) The mean Variance Accounted For (VAF) between subspaces for distinct social identities. (**F**) The VAF for each social subspace across 4 days. (**G**) Distributions of *p*-values for statistic comparison between the true VAF and the shuffled VAF (1000 times). Inserts: the proportions of shuffled data points above (red) and below (yellow) the chance level. Dashed lines indicate the α value at 0.05. See the Extended Table 1 for detailed statistic results.

Within the neural geometry framework, distinct stimuli are thought to engage in specific low-dimensional subspaces of population neuronal activity and form stable stimulus-specific representations(*42*). We, thus, tested whether each social demonstrator could be stably represented by a specific neural subspace. For example, for the demonstrator 1 (*D*_*1*_), we recorded differential spiking activities recorded in the test mice when socially interacting with the *D*_*1*_ housed at different arena locations (*D*_*1*_-*C*_*1*_, *D*_*1*_-*C*_*2*_) in the consecutive two tests (Fig. 3D), and after the joint SVD, these activity patterns formed an orthogonal basis to define a *D*_*1*_-specific subspace. Although similarities between different demonstrator subspaces were relatively low within a day (Variance Accounted For ratio, VAF < 63%) (Fig. 3E), similarities of the same demonstrator’s subspace across days were quite high with VAF > 90% (Fig. 3F), suggesting stable neural subspaces for encoding individual demonstrators across days. After the permutation test, we found that 83% of pairwise comparison was significantly stable (Fig. 3G). Moreover, we obtained similar results using the non-linear dimension reduction approach (fig. S4C to F). The subspaces derived from the social-related neurons were more stable than that from the non-social neurons across days (fig S5). Chemo-suppression of these responsive mPFC neurons during the multi-individual social test, which were genetically traced using the cFos-TRAP transgenic mice(*28, 43*), dramatically decreased the similarity or stability of demonstrator-specific subspace across days (when comparing the Clozapine *N*-oxide and saline groups; fig. S6).

Thus, these results suggest that social identities can be stably encoded by distinct low-dimensional geometric subspaces of population activity of active mPFC neurons for several days.

### Conditioned Social Learning Enhances Social Discrimination to Multi-individuals

Differential valences in social experience with different individuals can strongly affect strength of social discriminative memory(*1*). Based on this notion, we developed a “multi-individual social learning” paradigm, and it consisted of three stages (Fig. 4A): i, the pre-conditioning stage, where the test mouse underwent repeated social tests for 4 days to determine its initial social preferences rank to four unfamiliar demonstrators; ii, the social conditioning stage, in which each social duration of the test mouse with demonstrators ranking at the first (marked as *Aversive* demonstrator) and forth (marked as *Reward* demonstrator) positions was associated with aversive air puff and milk reward, respectively, for consecutive 8 days (Movie S3 to S4), while that with the remaining two demonstrators (marked as *Neutral 1* and *Neutral 2* demonstrator) had not any conditioning stimuli (as neutral), randomizing the conditioning orders and housing corners of individual demonstrators across days; and iii, the post-conditioning test: the test mice underwent the two sessions of multi-individual social test to determine a change of its social preference after the conditioned social learning (Movie S5 to S6). The behavioral results indicated that test mice did not have any overall preference to particular corner (Fig. 4B), while they showed ranks of their preference in social exploration to four different demonstrators in the pre-conditioning social test (Fig. 4C cyan). However, the reward and aversive social conditioning for 8 days substantially increased and decreased the test mice’s time durations of social interaction with the *Reward* demonstrator and the *Aversive* one, respectively, in the post-conditioning test, resulting in opposite shifts of preference towards to the *Reward* and *Aversive* demonstrators that were initially at the lowest and highest ranks, respectively (Fig. 4C purple). As a control, the test mice did not significantly change their social duration spent on other two *Neutral* demonstrator (Fig. 4C *Neutral* 1 & 2). This conditioning-induced social discrimination memory was largely unaffected when the test mice performed their post-conditioning test in completely dark environment (Fig 4D). However, we found that these transgenic mice lacking pheromone-sensing receptors (*Trpc2*-null)(*6, 44, 45*), the conditioning-induced changes of social preferences changes were diminished (Fig. 4E). These results strongly suggest that mice can behaviorally discriminate among at least three individuals, and such social discrimination function and memory for multiple individuals critically rely on chemosensory perceptions rather than visual perceptions of individual demonstrators.

**Fig. 4.**
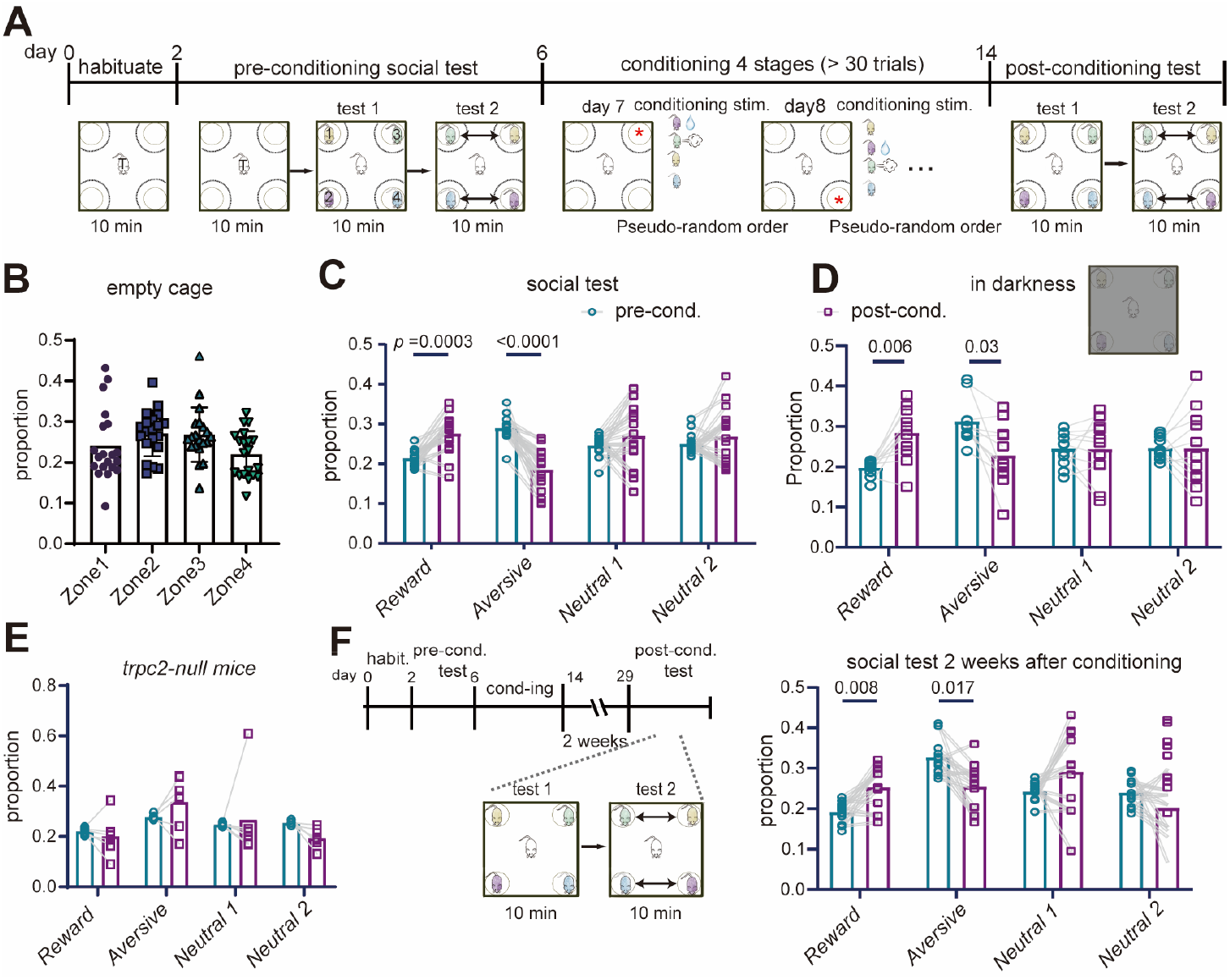
Recognition and memory of multi-social individuals tested by a social conditioning learning. (**A**) The social conditioning learning for multi-social individuals. The social conditioning is to associate milk reward, aversive air puff, and blank consequence with social investigations to 4 individual demonstrators in the test mice. The social conditionings with individual demonstrators were conducted at pseudo-random orders and at randomly selected corners across 8 days, followed by a post-condition multi-social test to evaluate the change in the rank of social preferences to individual demonstrators. (**B**) No spatial preference to individual locations of empty houses in the test mice, indicated by the proportions of exploration time spent on each corner. (**C**) Changes of the proportions of time for social investigations to 4 individual demonstrators pre-conditioning and post-conditioning the social conditionings shown in the (**A**). (**D**) Same as the **c** except the post-conditioning social test was conducted in darkness. (**E**) Same as the (**C**) except use of the *Trpc2-null* test mice. (**F**) The protocol (*left*) and results (*right*) of conditioned social discrimination memory tested 2 weeks after the social conditioning in the wild type mice. The *p* values were calculated by the Two-way ANOVA with Bonferroni’s multiple comparisons test (see detailed information in the Extended Table 1). *Habit*.: Habituation phase; *Pre-cond*.: Pre-conditioning social test; *Post-cond*.: Post-conditioning social test.

More interestingly, we found that the shifts of conditioning-induced social preference in test mice were still persisted in the test two weeks after the conditioned learning (Fig. 4F), implicating a form of long-term memory. This observation is distinct from these previously reported short-term social memory (lasting approximately 12 hr)(*13*). Moreover, in the post-conditioning multi-individual social tests, chemo-suppression of the responsive mPFC neurons, which were genetically traced using the cFos-TRAP transgenic mice in the conditioning phase, impaired the shifts of conditioning-induced social preference in test mice (fig. S7), supporting the key role of mPFC in social memory(*28, 43*).

### Reorganization of Neural Geometric Representations in Social Memory

Since we have identified distinct low-dimensional geometric subspaces of population mPFC activities as neural representations for individual demonstrators (Figs. 2 to 4), we further characterized how the conditioned social learning could alter these neural geometric representations. At the single-unit level, we observed trends of decreasing the proportion of *non-response* neurons in recorded mPFC units (Fig. 5A) and increasing of the proportions of *combinative* neuron after the social conditioning (Fig. 5B), in overall. It might implicate a possible recruitment of more neurons for this social discrimination test. At the population activity level, the SVM decoding accuracy in distinguishing among four demonstrators based on recorded population activity in the post-conditioning test was substantially higher (Fig. 5C), and more pronounced decoding enhancements were particularly for the *Reward* and *Aversive* demonstrator (Fig. 5D to G). These results suggest that social conditioning experience could enhance differential neural representations for individual identities, particularly for individuals associated with valence outcomes, a neural process accompanied with the animal’s behavioral shifts of its social preference rank.

**Fig. 5.**
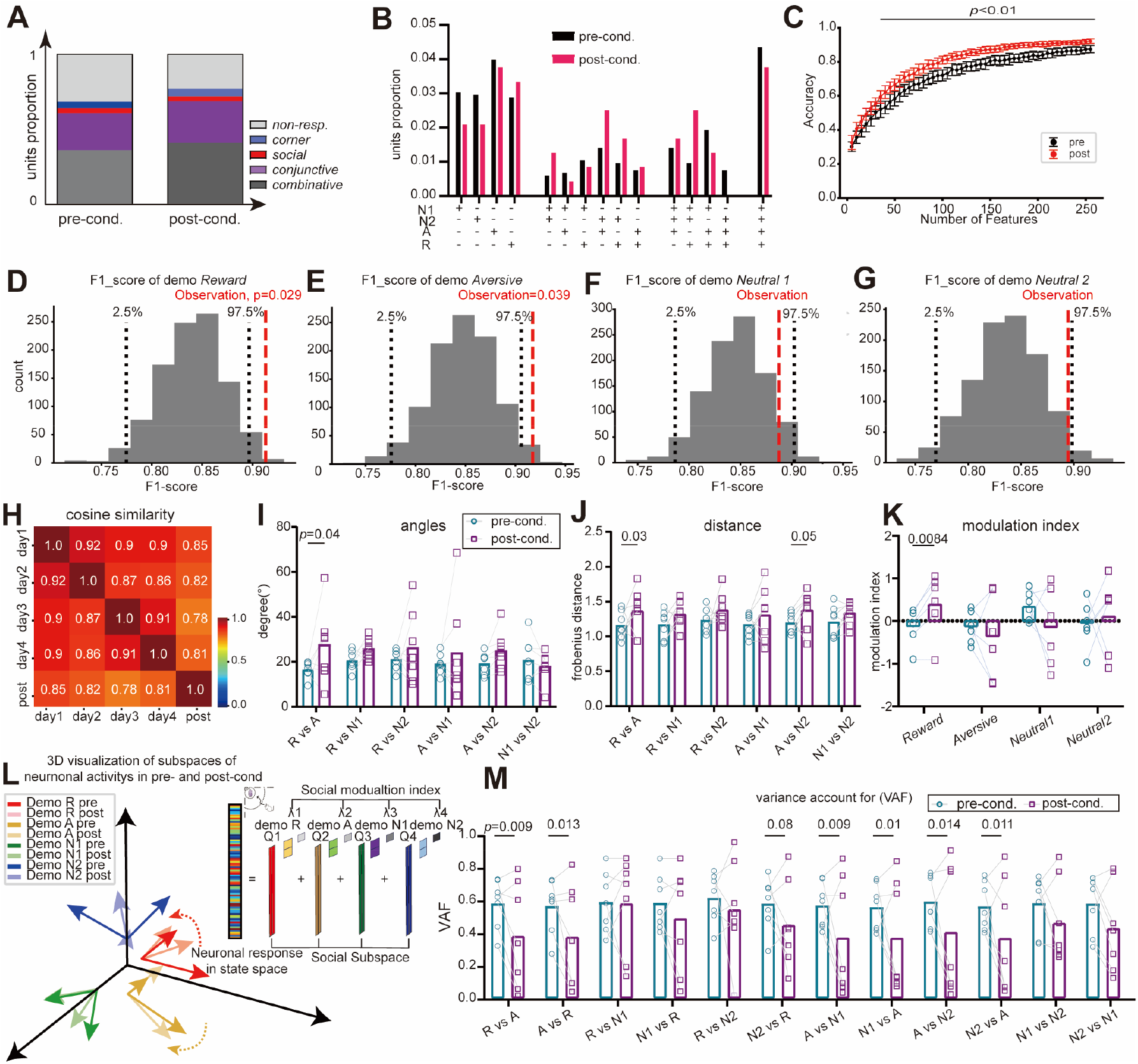
Social conditioning-induced selective enhancement of neural-activity representations of the reward demonstrator and Aversive demonstrator. (**A**) Changes of the proportions of different types of recorded neurons showing distinct selectivity in the multi-social tests before and after the social conditioning. Note that the proportion of *non-response* neurons significantly decreased (chi-square test, χ^2^ = 10.30,*p* = 0.036; Z_*non-resp*._ = 3.168, *p*_*non-resp*._ = 0.0076). *resp*.: response. (**B**) No significant changes in the proportions of *social* units encoding distinct demonstrators before and after the social conditioning (chi-square test, χ^2^ = 12.12, *p* = 0.67). (**C**) Consistent higher decoding accuracies in the SVM using the post-conditioning units than that using the pre-conditioning units, with increasing numbers of units (*: *p* < 0.01, t-test). R: *Reward* demonstrator; A: *Aversive* demonstrator; N1: *Neutral 1* demonstrator; N2: *Neutral 2* demonstrator. (**D** to **G**) F1 scores for population unit activity decoding of distinct *Reward, Aversive, Neutral 1* and *Neutral 2* demonstrators pre- and post-conditioning, indicating selective increases in the accuracy of decoding the *Reward* and *Aversive* demonstrators. (**H**) Stable population activity representations of social individuals across pre- and post-conditioning test days, indicated the cosine similarity after the joint SVD. (**I** to **J**) Changes of the angles and distances between distinct neural subspaces encoding *Reward, Aversive, Neutral 1* and *Neutral 2* demonstrators, respectively, pre- and post-conditioning. Note the significantly-increased angle and distance between the *Reward* and *Aversive* demonstrator subspaces (Two-way ANOVA with Tukey’s multiple comparisons test). (**K**) Changes of the modulation index of neural subspace representations for individual demonstrators pre- and post-conditioning, indicating a selective enhanced intensity in the *Reward* demonstrator representation (Two-way ANOVA with Tukey’s test). (**L**) A schematic illustration of changes in neural subspace geometries for encoding distinct demonstrators based on angle and distance metrics (as shown panels **I** to **J**). (**M**) VAF for distinct demonstrator subspaces pre- and post-conditioning, showing specific reduction of VAF for Reward and Aversive demonstrator, respectively (Two-way ANOVA with Tukey’s test, see detailed information in the Extended Table 1).

Furthermore, we found that the mPFC population activity after the social conditioning experience could still be embedded in a similar low-dimensional space when compared with those from the pre-conditioning tests (Fig. 5H bottom rows). However, we identified several key changes within demonstrator-specific subspaces as follows (Fig. 5I to M). We noted that the angle and distance between the *Reward* and *Aversive* subspaces significantly increased at the post-conditioning stage, suggesting an enhanced separation between the neural subspace for these valence-opposed demonstrators (Fig. 5 I to J). It was further evident by a decrease in the VAF ratio (Fig. 5M). Secondly, the modulation index, which reflects the magnitude of population-level responses to a given stimulus(*32, 36*), was dramatically elevated in the neural subspace for the *Reward* demonstrator, indicating a stronger engagement of neurons encoding this specific individual (Fig. 5K). It was also consistent with the above observed improvement in population-level decoding accuracy (Fig. 5D). Meanwhile, enhanced angular separations and reduced shared variance (VAF) were found between the *Aversive* and *Neutral* subspaces (Fig. 5M), suggesting that discriminability among these subspace representations was likewise enhanced as well.

In summary, these results demonstrate that social conditioning experience effectively drives specific alterations in respective social-identity subspaces, and selectively enhance separations of subspaces for social-behaviorally salient individuals along with moderate refinements of representations among others. These reorganizations of neural geometric representations in mPFC neurons may promote discrimination among interacted individuals.

### Evolution of Conditioning Responsive Neurons to Social Neurons during Learning

A previous study suggested that learning-induced changes in neural geometric properties could contribute to the improved ability of performing certain behaviors(*42*). We, thus, attempted to elucidate how the conditioned social stimulus or experience would reshape neural geometric representations of individual identities in mPFC neurons. We first analyzed mPFC neuronal spiking responses during the conditioning period, using the Elastic-Net regression (L1 + L2 regularization) method (fig. S8A to B, and ref. (*29*)) to capture encoding of specific event features (e.g. the demonstrator, positive milk reward or aversive air puff stimuli) by spiking activities. As the demonstrator was housed in a fixed corner during daily conditioning, we excluded the spatial corner features in the latter analysis. The analysis indicated that during social conditioning, spiking rates of responsive mPFC neurons were modulated exclusively by a demonstrator (Fig. 6A; *social* neurons) or conditioning reward /aversive stimuli (Fig. 6B; *conditioning* neurons), or jointly by both the demonstrator and its associated conditioning stimuli (Fig. 6C; *joint-response* neurons). There existed *non-response* neurons showing no changes in their spiking rates during the conditioning (Fig. 6D). It was worth noting that the daily proportions of *joint-response* neurons were increased within the 8 days of conditioning, while that of other two types were not significantly altered (Fig. 6E), implicating a potential learning-induced shift of neuronal encoding to integrated features of social individual and its associated valence. On the contrary, proportions of demonstrator-specific *social* neurons and the neural space similarity of social representations in the joint SVD were kept relatively stable, respectively, throughout the conditioning period (Fig. 6F to G).

**Fig. 6.**
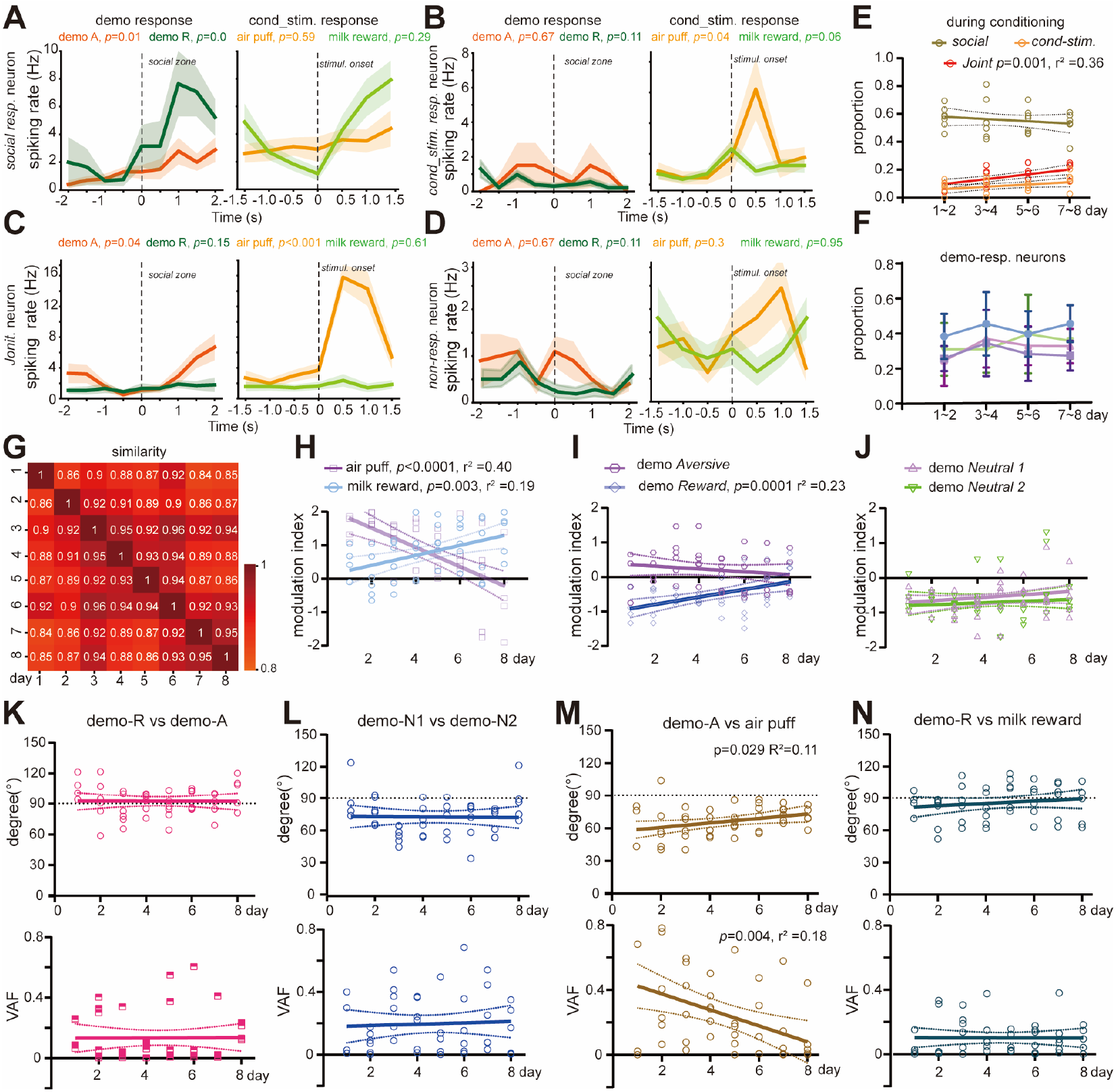
Preferential shifts of neural representations toward the Reward demonstrator during social conditioning. (**A** to **D**), Representative *social response* unit (**A**), *conditioning response* unit (**B**), *joint-response* (social and relative conditioning stimulus) unit (**C**) and *non-response* unit (**D**), showing differential changes in spiking rates (binned with 500 ms) before and after the onset of entering the social zone (*left*) or receiving conditioning stimuli (*right*), respectively, in the social conditioning. (**E**) Changes of the proportions of these three types of responsive units across the 8 conditioning days. Note a trend of increasing the proportion of *joint-response* units with the conditioning days (simple linear regression). (**F**) Similar as **e** except proportions of demonstrator-specific encoding units were analyzed. (**G**) Cosine similarity of population activity representations in the joint SVD among the 8 days of social conditioning. (**H** to **J**), Opposite changes of the modulation index of neural subspace representations for reward and aversive condition stimulus (**H**), or *Reward* and *Aversive* demonstrator subspace (**I**), while there is no significant change for *Neutral 1* and *Neutral 2* subspace (**J**) during the 8 days of social conditioning. (**K** to **N**), Comparisons of dynamic change of the angles (*top*) and VAF (*bottom*) between paired individual neural subspaces for distinct demonstrators or conditioning stimulus. Note that subspaces for the *Aversive* demonstrator and the air-puff stimuli become more separated by increasing the angle and reducing VAF simultaneously during the 8 days of social conditioning.

Furthermore, we noted that the population activity modulation indices for different conditioning-related feature representations showed differential changes across conditioning days. In the early conditioning days, modulations by the aversive air-puff stimuli and its associated *Aversive* demonstrator were higher, but decreased dramatically with condition-day continuing (Fig. 6H to I purple). On the contrast, modulations by the reward stimuli and its associated *Reward* demonstrator consistently increased with conditioning and surpassed the level of aversive stimuli-related modulations during conditioning days (Fig. 6H to I light blue). However, no significant changes were observed in the modulations by the *Neutral* 1 or *Neutral* 2 throughout the conditioning period (Fig. 6J). These results implied a strategic evolution from avoiding punishment to pursuing reward in the animal mind during conditioned social learning.

More importantly, several key evolutions of demonstrator or conditioning stimuli-specific low-dimensional neural subspace were observed over the conditioning period. We noted that the orthogonal angles and tiny VAF values between the *Reward* and *Aversive* demonstrator subspaces consistently kept over conditioning (Fig. 6K), while subspaces for the *Neutral 1* and *Neutral 2* demonstrators showed less angles and relatively higher VAF (Fig. 6L). Meanwhile, the subspace for the *Aversive* demonstrator also gradually shifts separative to that for *Neutral 1* and *Neutral 2* demonstrators over the conditioning (fig. S8C to F), suggesting that a growing separation was observed between the *Aversive* demonstrator and the other. However, over the conditioning progress, subspaces for the *Aversive* demonstrator and air-puff stimuli gradually converged toward orthogonality, accompanied with their shared variance decreasing (Fig. 6M), suggesting that the animal gradually learn to dissociate the aversive stimulus from the social individual. Such evolution was not observed in the subspaces between the *Reward* demonstrator and the reward stimuli (Fig. 6N).

Together, these results demonstrate that the conditioned social learning not only drives continuous emergences of spiking neurons encoding integrated social and valence information, but also selectively reshape geometric angles and distances among respective neural representational subspaces for various demonstrators and conditional stimuli over time. All these evolutions of mPFC population activity may support a strategic reorientation of social preference to respective demonstrator during social learning.

## Discussion

In this study, we employed a novel one-*versus*-four social discrimination paradigm combined with *in vivo* electrophysiological recordings, demonstrating that mice can behaviorally discriminate among multiple social individuals. We found that only a small fraction of neurons in the mouse mPFC exhibited selective responses to specific social individuals, whereas the majority displayed multi-selectivity properties (Fig. 1). Furthermore, our data revealed that social representations in this region are highly nonlinear (Fig. 2). Using a newly developed algorithmic method, we uncovered that the mPFC stably represents different social individuals within distinct neural subspaces at the population level (Fig. 3). These subspaces undergo experience-dependent angular shifts following social encounters, which in turn modulate subsequent social preference (Fig. 4-6). And mPFC played the key role in social subspace stability and social preference memory (fig. S5 to S7). These findings provided a computational framework for understanding complex social cognition.

Most previous studies on social memory have widely used the traditional three-chamber test(*7-13*), which exploits the preference of mice to spend more time on investigating a novel conspecific than a familiar one, thereby providing a framework for studying the formation of social memory. However, numerous studies have revealed that neurons in the mPFC and hippocampus often exhibit multi-selectivity, a property thought to support the storage of more memory(*14, 15, 23, 29, 30*). Consequently, relying solely on dichotomous discrimination tasks to assign neuronal selectivity may be overly simplistic. Although several studies have examined multi-individual recognition in mice and reported social memory coding in dorsal CA1(*16*), this region is predominantly associated with spatial encoding(*46*) and an explicit description of social-specific representations has remained unknown. Our study with *in vivo* recordings from freely moving test mice has demonstrated that mice can discriminate among multiple social individuals (Fig. 1-5), and at population neuron level, social stimuli representations are highly nonlinear in the mPFC (Fig. 2). Moreover, with the conditioned social learning, mice gradually refined their discrimination among demonstrators that are associated with positive, negative or neural experiences, respectively. It is noteworthy that Hopfield Recurrent Neural network simulations, in which implement the recurrent neural connectivity and Hebbian plasticity to generate stable attractors for distinct representations, have strongly suggested high capacity and low abstraction properties of population level representations of social individuals in the mPFC (*36-39*). Our experimental observations (Fig. 2E to G) agree with this notion and are reminiscent of the neural dynamics in Hopfield-type networks. Although the mPFC displays high representational capacity, it is a high-order region positioned toward relatively later stages in the neural processing hierarchy. Thus, it is very likely that information to the mPFC is already substantially abstracted from these upstream areas(*47*), and thus high encoding capacity of mPFC neurons may reflect a storage of such well-processed information. Disentangling degrees of information abstraction to the mPFC demands brain-wide recordings across multiple nodes of the social processing network(*47*).

Electrode drifting has been known to impede a longitudinal tracking of neural activities from a same population across days(*48*). It limits a robust analysis to changing neural presentations for learning or memory over time. At neuronal population activity level, In this study, we involved in the framework of neural manifolds(*42*). Previous studies have established that the manifold analysis has been implemented to delineate neural representation of specific actions or cognitive processes across sessions(*49*). Moreover, neural activity geometries imbedded these manifolds further offer meaningful comparisons across days(*33*), individuals(*33, 50*) and even species(*51*). In the present study, we have combined neural geometry analysis, linear regression and a joint SVD-CCA approach to align population activity of different days into the same low-dimensional subspaces and compare different among social subspaces(*31, 33*). With this approach, we have revealed that distinct social individuals are encoded within separate, highly stable low-dimensional subspaces in the mPFC (Fig. 3). Notably, our results also suggest that positive or aversive social experience triggers substantial changes in neural geometry, a process that may underlie the shifts of social preference rank for different demonstrator following the conditioned social learning.

The “concept cells” in the human medial temporal lobe was proposed by Quiroga and colleagues, where neurons responded selectively to specific individuals (e.g. the actor Jennifer Aniston)(*52*). This study has spurred identifications of analogous “engram cells” in the ventral hippocampal CA1(*16, 19, 20*) and CA2(*10, 18*) as well as the mPFC(*6, 8, 9, 12, 21, 47*) in rodents. However, how multiple social individuals are dynamics represented in these circuits and how changes of the representations regulate social actions or preference remain unresolved. The present study suggests that mPFC neurons progressively integrate the valence information in interactions with related social demonstrator, and in the latter process we observed a significant increase of the proportion of neurons exhibiting conjunctive tuning to both social identity and associated valence (Fig. 6). Moreover, neural representations of individuals associated with positive (reward) valence is selectively strengthened progressively along with social conditioning, whereas those linked to negative (aversive) valence are weakened, leading to sharpen distinction between reward and aversive demonstrators in neural geometry. Together, these dynamic changes in population-level coding can serve as a mechanistic substrate for experience-dependent shifts of the social preference among multiple individuals.

## Acknowledgments

We would thank Dr. Qian Li (SJTU) for providing the Trpc2-null mice and Dr. Haitao Wu (BIBMS) for supporting whole-brain section imaging.

## Author contributions

Conceptualization: X-h.Z., T-y.L.

Funding acquisition: X-h.Z.

Methodology: X-h.Z., T-y.L.

Experimental design: X-h.Z., T-y.L.

Experimentation and analysis: T-y.L., X-z.H

Visualization: X-h.Z., T-y.L.

Writing – original draft: X-h.Z., T-y.L.

Writing – review & editing: X-h.Z., T-y.L.

## Data, code, and materials availability

The preprocessed data and code of this paper is available on GitHub (https://github.com/Tianyulied/Neural-geometric-representations-of-social-memory-for-multi-individuals-in-medial-prefrontal-cortex) and on Zenodo (https://zenodo.org/uploads/18538407). The raw data (electrophysiological recordings and video files) are not publicly available due to their substantial size. They are available from the corresponding author upon reasonable request.

## Supplementary Materials

Materials and Methods

Supplementary Text

Figs. S1 to S8

Tables S1

References (*53*–*56*)

Movies S1 to S6

## Materials and Methods

A portion of code were generated with the assistance of Bing Copilot and Deepseek-R1. All AI-generated code was thoroughly reviewed, tested for accuracy, and modified by the authors to ensure it meets the standards required for the analyses presented in this work. The authors assume full responsibility for the integrity of the code and the results.

### Animals

Adult wild type (WT) C57BL/6J mice (at postnatal 3-6 months, from the Beijing Vital River Laboratory Animal Technology Company.) were used in the most of animal behavioral tests and *in vivo* recording experiments. In several sets of experiments, two additional transgenic lines, the *Trpc2* knock-out mice (021208; RRID:IMSR_JAX:021208) and the cFos-CreER (Jax. No. 021882) were used as the test mice as well(*43*). Only male mice were used in this study. In the behavioral tasks, one WT or transgenic mouse was used as the test mouse, while other four mice, at matching ages, were selected as the social stimulus or demonstrator mice in the multi-individual social test from other different cages. During the conditioning phase, the test mouse was regularly housed with its littermates, and after being implanted with a recording chamber, it was then housed alone in order to protect the implanted electrode. All mice were raised on a revered 12 h light (18:00–6:00)/dark (6:00-18:00) cycle, under an environmental condition with 23∼25 °C ambient temperature and 40-60 % relative humidity. The protocols of animal surgery and experiments were approved by the Animal Care and Use Committee (IACUC) of State Key Laboratory of Cognitive Neuroscience & Learning at Beijing Normal University (reference NO.: IACUC-BNUNKLCNL-2021-17).

### Animal Surgeries, Electrode-array Implantation and Viral Injection

Mice were anesthetized using isoflurane (4% for induction, 0.5–1.5% for maintenance) and tightly positioned on a stereotaxic apparatus (RMD, China). A custom-built micro-drive equipped with a 32-channel micro-wire array (KD-MWA, Kedou Brain-Computer Technology, China) was implanted in the medial prefrontal cortex (mPFC) at one hemisphere. Stereotaxic coordinates (relative to the Bregma) were: +1.9 mm AP, + 0.25 mm ML, and –0.8 to –1.0 mm DV from the cortical surface, to make sure all wire in PL or IL area (Extended Data Fig. 1). Then the micro-drive assembly was affixed to the skull using three small stainless-steel screws (M1, 1mm), cyanoacrylate adhesive and dental acrylic. After the implantation, animals were individually housed to recover on a heating pad until fully ambulatory.

For viral injection, the cFos-CreER mice (at postnatal 8-9 weeks) were fixed on a stereotaxic device under the same anesthesia condition. The skull was exposed under antiseptic conditions and a small craniotomy was made with a thin drill over the prefrontal cortex. (typical coordinates: 1.9 mm anterior to Bregma; ± 0.3 mm lateral to the midline). An electronic injector (RWD, China) attached to the stereotaxic devise was used to infuse ∼300 nL viral solution (*pAAV-EF1α-DIO-hM4D(Gi)-mcherry-WPRE*, 1.12 x 10^13^ particles per ml, Obio Technology, Shanghai) into the target cortical region through a borosilicate glass micropipette (tip size, 10 μm), at a delivery rate of 20 nL/min (fig. S6 to S7). At the end of injection, the micropipette was held in the place for about 10 min before retraction. The mice were then placed in a stable-temperature incubator to recover from anesthesia, and after a full recovery they were returned to their home cages.

### Drug injections

In cFos-CreER mice experiments, tamoxifen (Sigma) was dissolved in corn oil (Sigma) at 37°C overnight to a concentration of 20 mg/mL and subsequently administered to cFos-CreER mice via oral gavage at a dose of 150 mg/kg 24 hours prior to behavioral testing. Tamoxifen solution was stored at 4°C for no more than 24 hours before use.

Clozapine *N*-oxide (CNO, MCE) was dissolved in DMSO (Sigma) at 20 mM and stored at −20°C. Before experiments, stock CNO was diluted in saline to the desired concentration, 0.4 mg/mL for hM4Di. The same amount of DMSO was diluted in saline as vehicle control. Fresh CNO was injected i.p. at 4 mg/kg 30 minutes before behavioral testing.

### Confocal imaging

Animals were either perfused or drop fixed with 4% paraformaldehyde (PFA, Sigma) in PBS. Coronal sections were cut at 80 μm thickness for checking micro-wire array tip and 40μm thickness for others. Slices were washed in PBS for 10 minutes 3 times and mounted with DAPI Fluoromount-G (Southern Biotech). Micro-wire array images were acquired using an Olympus BX53 System microscope. And other images were acquired using a Nikon A1R confocal microscope (Japan) with a Plan-Apo 20X objective (0.75 N.A., Nikon, Japan.

### Multi-individual Social Test

#### Apparatus of Multi-individual Social Test

A custom-built wooden square arena, 50 cm (length) × 50 cm (width) × 50 cm (height), was used for the behavioral social tasks, in which a test mouse made social interaction and recognition with other four demonstrator counterparts (Figure 1A). The demonstrator mice were housed in a wire pencil cup (8 cm diameter and 10 cm heigh) and positioned at four corners individually in the arena. A semi-circle area with 3.5 cm away from the cup’s circle edge was defined as the social investigation zone at each corner(*53*), and its outer edge was installed with six infrared photoelectric sensor to detect the time of entering and leaving the zone precisely. The arena was lit evenly during the test, and animal’s behaviors in the arena were recorded by a digital camera (IRAYPLE A7300MU90; 60 fps, 2048×1536 pixels) mounted 80 cm above the center of the arena. In some experiments, two additional cameras were used to record from the arena sides. Recorded videos were analyzed by the DeepLabCut software(*54*) (version 2.3.9, https://github.com/DeepLabCut/DeepLabCut?tab=readme-ov-file), in which the test mouse’s positions were tracked and its defined social investigation actions (e.g. sniffing and touching the demonstrator mice in the social zone) as well as other non-social behaviors (locomotion and stationary manners) were accordingly labelled and categorized for further analyzed.

Both test and demonstrator mice were separately habituated to the arena for 2–3 days prior to experimentation. Our social behavioral paradigm comprised three phases: the social preference rank test (pre-conditioning phase), the social conditioning, and the post-conditioning social preference rank test. In above phases, all controls to video-recording, social investigation detection, conditioning stimuli delivery and time alignments of electrophysiology recording were conducted by an Arduino-based finite-state machine (BPod r0.5, Sanworks) and our own custom MATLAB scripts (MathWorks, version R2023b).

#### Social Preference Rank Test (pre-conditioning phase)

After 2- or 3-day habituation to the testing arena, the test mice encountered four housed demonstrator mice fixed at 4 corners for the first time. The test mouse was allowed to freely explore the arena and make social investigations to individual demonstrator mice for 10 min. Its time spent within individual demonstrator’s social zones were calculated. To minimalize potential biases to specific corners or demonstrator in the test mouse, we set two 10-min test sessions, in which the corners of housing all demonstrators were swapped each other (as shown by the schematic diagram in the Figure 1A) in each day. For each test mice, its proportions of social investigation time spent on four individuals were averaged from the two sessions across 4 days to obtain the social preference rank.

#### Social Conditioning Protocol

Based on the initial social preference rank order for a test mouse, we then set the positive milk reward and aversive air puff stimuli paired with social interaction when the test mice were socially investigating these two demonstrators that resized the first and the last ranks, respectively. Thus, the latter two demonstrators were designating as the *Aversive* demonstrator and *Reward* demonstrator, respectively. Meanwhile, social investigation of the test mice to the rest two demonstrators yielded no any outcome, and similarly these two demonstrators were designating as the *Neutral 1* and *Neutral 2*.

Each test mouse was conditioned during its social interactions with *Reward, Aversive, Neutral 1* or *Neutral 2* demonstrators separately in the same arena 8 days (Figure 4A). Details of social conditioning stimuli were as follows:

When the test mouse made a social investigation to individual housed demonstrator ≥ 3 seconds in the social zone, a specific outcome was triggered based on the designated demonstrator identity:

- ***Reward* demonstrator**: 2 drops of sweet milk delivered outside the social zone to promote continued social exploration (Movie S3).
- ***Aversive* demonstrator**: An air puff was administered to discourage interaction (Movie S4).
- ***Neutral* demonstrators**: No stimulus was delivered.

A daily conditioning session comprised ≥ 30 trial or lasted up to 20 min for individual demonstrators, respectively. Corner positions and conditioning trial sequences for different demonstrators were daily randomized to avoid possible preference to specific corner or conditioning order in the test mice (Figure 4A).

#### Post-conditioning Multi-individual Social Test

After conditioning, mice underwent two consecutive social tests wherein all demonstrator positions were changed between sessions to control for spatial and novelty effects. Changes in social preference were evaluated using Two-way ANOVA. For statistical details, see table S1. ***in vivo* Electrophysiology in Behaving Mice**

A 4×8 array of KD-MWA micro wires (Nitinol, 25μm, impedance 100kΩ-300kΩ) was implanted into the mPFC of the test mouse. A wire was inserted above V1/2 area as a common reference. The microwire array was connected to a 32-channel head-stage, and electric signals were recorded by the CerePlex-Direct data acquisition system (Blackrock Microsystems, USA). Neuronal spiking signals from extracellular recording were digitized at 40 kHz, bandpass-filtered (250 Hz - 8 kHz) and acquired by a computer for further analysis. The *in vivo* recording was made while mice freely explored the environments and housed demonstrator mice in arena for 10 minutes. A commutator was used to reduce the cable torques on the head-stage and minimalize any possible restraint on mouse’s free movements. Time of recording neural signals was synchronized with that of video-recording behaviors.

We used the KiloSort 2.5 (https://github.com/MouseLand/Kilosort) to sort raw spike waveforms into single units (presumptive neurons)(*55*), using the following parameters: Nblock = 0 (optimized for micro-wire arrays), detection threshold = 9, lambda = 10, and AUC threshold = 0.7. Putative units identified by the KiloSort as “good” or “mua” (multi-unit activity) were subsequently manually curated in the Phy 2.0 (https://github.com/kwikteam/phy), based on waveform. Units with firing rates <0.3 Hz were omitted. Final unit inclusion was based on the following quality metrics, computed via the SpikeInterface (https://github.com/SpikeInterface/spikeinterface): inter-spike interval violation ratio < 0.5, presence ratio > 0.95 per session, amplitude cutoff < 0.1, and signal-to-noise ratio (SNR) > 2.

### Linear regression model

To identify task variables modulating neural activity during pre- and post-conditioning social tests, we employed a regularized linear regression model. Eight one-hot encoded task variables *S*_demo,loc_ were defined, corresponding to each combination of demonstrator identity— *Neutral 1, Neutral 2, Aversive*, or *Reward*—and its spatial location across the two test sessions (4 demonstrators×2 sessions = 8 combinative variable).

The firing rate *y*_*i*_ (binned into 250-ms interval) of the ***i***th neuron was modeled as a linear combination of these task variables:

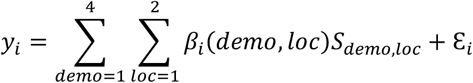

Where *β_i_*(*demo, loc*) denotes the regression coefficient for each demonstrator-location predictor, and ε*_i_* represents the residual error. To mitigate overfitting and perform feature selection, we applied L1 (Lasso) regularization. The regularization strength was optimized via 10-fold cross-validation (*31*).

To identify neurons significantly encoding a specific predictor, we compared full and reduced models using a cross-validated variance-explained criterion. For each neuron, we iteratively refit the model on 90% of the data (10-fold cross-validation), each time omitting one predictor of interest. A significant increase in the mean squared error (MSE) of the residuals upon predictor removal indicated that the neuron encoded that variable.

Neuronal classification was based on spiking response characteristics during the pre- or post-conditioning social test:

- ***Social* neurons**: significantly responsive to the same demonstrator across different locations.
- ***Corner* neurons**: significantly responsive to the same location across different demonstrators.
- ***Combinative* neurons**: significantly responsive to a specific demonstrator-location combination.
- ***Conjunctive* neurons**: met criteria for both *social*- and *corner*-response units.

All *social, conjunctive*, and *combinative* neurons were classified as social-related neurons; the remainder were categorized as non-social neurons. During the conditioning phase, where all demonstrators occupied the same location, spatial confounds were controlled, and encoding was assessed solely based on the significance of residual MSE increase.

To assess how task variables influenced neural activity during the conditioning phase, we employed a multivariable linear regression model similar to that used in the social tests, but with an expanded set of predictors. Task variables included demonstrator identity (*Neutral* 1, *Neutral* 2, *Aversive*, or *Reward*) and conditioning type (Milk reward or Air puff). Since Milk reward and Air puff stimuli occurred as discrete events but could elicit sustained neural responses, we utilized Elastic Net regularization—a hybrid approach combining L1 (Lasso) and L2 (Ridge) penalties. This method simultaneously performs feature selection (via Lasso) and models temporal persistence in neural activity (via Ridge, fig. S8A)(*29*). The regularization strength was optimized via 10-fold cross-validation. Neuronal classification was based on spiking response characteristics during the conditioning phase:

- ***Social* neurons**: significantly responsive to the social demonstrator.
- ***Condition* neurons**: significantly responsive to Milk reward or Air puff.
- ***Joint-response* neurons**: significantly responsive to Milk reward with demonstrator *Reward*, or to Air puff with demonstrator *Aversive*.

### Population decoding

To determine whether population activity patterns in the mPFC encode discriminable social information and to characterize the linearity of such representations, we performed population decoding using the support vector machines (SVMs). We trained the multi-class SVMs with both linear and radial basis function (rbf, non-linear) kernels to classify neural population activity into one of the four demonstrator identities.

For each recording session, the SVMs were trained and evaluated independently at each social interaction time point using 5-fold cross-validation. Model performance was quantified both by decoding accuracy (proportion of correct classifications) and the F1 score. The F1 score, the harmonic mean of precision and recall, provided a robust measure of classifier performance, especially under class imbalance, and offered a more comprehensive evaluation of the model’s ability to discriminate neural activity patterns associated with different social stimuli.

The F1 score was calculated as:

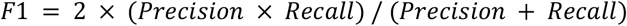

where *Precision*: The proportion of predicted positive samples that are actually positive.

∘ *Precision* = TP / (TP + FP)
∘ TP (True Positive): Correctly predicted positive samples
∘ FP (False Positive): Incorrectly predicted positive samples

- *Recall*: The proportion of actual positive samples that are correctly identified as positive.
  - *Recall* = TP / (TP + FN)
  - FN (False Negative): Incorrectly predicted negative samples

To assess the statistical significance of the decoder performance, we performed a non-parametric cluster-based permutation test (1000 times).

### Quantifying Representational Geometry with CCGP and Shattering Dimensionality

To assess the geometry of neural population representations and their support for abstraction versus specific encoding, we employed two complementary metrics derived from population decoding analysis: the Cross-Condition Generalization Performance (CCGP) and the Shattering Dimensionality (SD) (*36, 37*). Our experimental design generated 8 unique conditions, defined by the specific Demonstrator-in-Corner pairing (e.g., Demo 1 in Corner 1). This structure allows us to define variables of interest, such as corner or demonstrator identity, which are disentangled from absolute spatial location due to the full positional switched between tests.

#### Cross-Condition Generalization Performance (CCGP)

CCGP was used to quantify whether a given variable (e.g., corner or demonstrator identity) was encoded in an abstract format that generalizes across specific, untrained sensory combinations. For a target binary variable: We trained a linear support vector machine (SVM) decoder to classify the variable using neural data from only a subset of the experimental conditions (for example, data from Demo 1 in Corner 1 in test 1 and from Demo 1 in Corner 2 in test 2). The decoder’s performance was then evaluated on held-out testing conditions that belonged to the same variable categories but used the remaining, untrained corner-demonstrator pairings (for example, data from Demo 2 in Corner 2 in test 1 and from Demo 1 in Corner 1 in test 2; to test Corner CCGP). The CCGP value was calculated as the average test performance across all possible balanced splits of conditions into training and testing sets.

#### XOR Decoding Accuracy

To probe for more complex, non-linear interactions in the representation (e.g., whether the combination of corner and social demonstrator is encoded), we assessed the decoding accuracy for an exclusive OR (XOR) variable (trial decoding) (*36, 37*). A linear SVM was trained and tested using standard cross-validation across all trials for the relevant conditions. High decoding accuracy for such an XOR indicates that the neural population activity resides in a higher-dimensional space where non-linearly separable categorical boundaries can be resolved by a linear readout, a signature of complex mixed selectivity.

#### Shattering Dimensionality (SD)

SD measures the expressive capacity and linear separability of the neural population code across the entire space of possible binary classifications(*36, 37*). We enumerated all 28 unique balanced dichotomies. For each of these dichotomies, we computed the mean classification accuracy of a linear SVM decoder using standard cross-validation. The Shattering Dimensionality (SD) is defined as the average of these accuracy values. An SD close to 1 indicates a high-dimensional, linearly-rich representation where almost any arbitrary grouping of conditions can be decoded, signifying maximal flexibility for downstream computation. An SD near 0.5 indicates a representation that does not support linear separation of most variables.

### Hopfield Network Simulations of the Nonlinear Patterns

To model neural representations with varying degrees of mixed selectivity, we generated four binary memory patterns corresponding to the four experimental conditions: two social identities (Demo 1 and Demo 2) encountered at two corners (Corner 1 and Corner 2). The generation process was designed to produce patterns with controllable levels of nonlinear mixed selectivity (*36-39*).

We first generated 6 independent basis vectors *b*_l_, *b*_2_,…, *b*_6_, where each *b_i_* ∈ *R*^*N*^ was drawn from a standard normal distribution and subsequently orthogonalized using the Gram-Schmidt process:

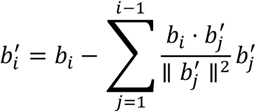

where *N = 200* represents the number of neurons in the network. Each orthogonalized vector was then normalized to unit length:

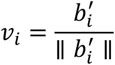

The six orthogonal basis vectors were assigned specific semantic roles:

- *v*_l_: Social identity A encoding vector
- *v*_2_: Social identity B encoding vector
- *v*_3_: Left position encoding vector
- *v*_4_: Right position encoding vector
- *v*_S_: Identity-position interaction term
- *v*_6_: Random component for increasing dimensionality

For a given nonlinearity level *λ* ∈ [0,1], each pattern *p_k_* was constructed as a weighted combination of these basis vectors:

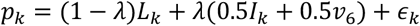

where:

1. **Linear component *L*_*K*_**:
  - For A-left: *L*_l_ = *v*_l_ + *v*_3_
  - For A-right: *L*_2_ = *v*_l_ + *v*_4_
  - For B-left: *L*_3_ = *v*_2_ + *v*_3_
  - For B-right: *L*_4_ = *v*_2_ + *v*_4_
2. **Nonlinear interaction term *I*_*K*_**:
  - For conditions with identity A: *I_k_* = *v*_5_
  - For conditions with identity B: *I_k_* =−*v*_5_ (sign inversion to enhance discriminability)
3. **Pattern-specific noise** 𝝐_*K*_:

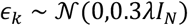

where *I_N_* is the *N* × *N* identity matrix.

The continuous-valued patterns were converted to binary states appropriate for Hopfield networks through a sign function:

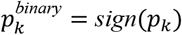

where the sign function operates element-wise, mapping positive values to +1 and non-positive values to -1.

Finally, each pattern was normalized to unit length:

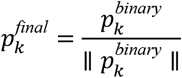

The nonlinearity parameter *λ* controls the trade-off between linear separability and representational dimensionality:

- *λ* = 0: Purely linear representations where identity and position are encoded in orthogonal subspaces, resulting in high cross-condition generalization performance (CCGP) but limited memory capacity.
- *λ* = 1: Highly nonlinear representations with strong mixed selectivity, where each specific identity-position combination has a unique encoding pattern. This yields low CCGP but high XOR decoding accuracy, indicating increased memory storage capacity.

For each set of generated patterns *p*_l_, *p*_2_, *p*_3_, *p*_4_, we trained a Hopfield network using the standard Hebbian learning rule:

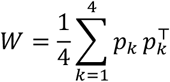

with the constraint of no self-connections (*w*_{*ii*} 0 *for ii* 1, …, *N*).

### CEBRA for Cross-Session Neural Embedding

To uncover latent neural dynamics and enable alignment of population activity across multiple days, we applied the CEBRA (Consistent EmBeddings of High-dimensional Recordings using Auxiliary variables; https://cebra.ai), a machine learning method designed for nonlinear dimensionality reduction of time-series data(*41*). CEBRA learns a mapping from high-dimensional neural activity into a low-dimensional embedding space that preserves temporal, behavioral, or neural correlation structure(*41*).

We used CEBRA in discrete, multi-session mode to reduce dimension of neuronal spike data binned into 250-ms intervals and aligned the test mouse social state across session with the following configuration: temperature_mode = “auto”, learning_rate = 3e−4, max_iterations = 10000, time_offsets = 10, output_dimension = 3, distance = ‘cosine’, and conditional = ‘time_delta’. This set of parameters was chosen to prioritize temporal consistency across sessions while capturing discriminative features related to social stimuli.

The resulted embeddings were then used to quantify representational stability and similarity across days. We computed session-pairwise similarity using the adaptive Hausdorff distance(*56*), which provides a robust metric for comparing the geometry between two sets of neural trajectories in the embedded space. This approach allowed us to assess whether social representations occupied consistent subspaces across days and conditioning phases.

### Joint SVD Projection Method for Cross-Dataset Neural Response Analysis

To enable consistent comparison of neural representations across day, we developed a unified dimensionality reduction framework based on joint Singular Value Decomposition (SVD). This method identifies shared neural response patterns while preserving dataset-specific variance structures through subspace orthogonalization(*33*).

Let 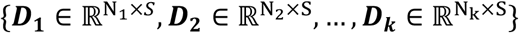 denote *k* neural response datasets, where *N*_*k*_ represents the number of neurons in dataset ***k*** and ***S*** denotes the number of stimulus conditions (i.e., combinations of demonstrator and location, the total amount = 8). Each entry in ***D***_**k**_ corresponds to a centered regression coefficient *β_i_*(*demo, loc*) for a given neuron and stimulus.

#### 1. Global Matrix Construction

All datasets are vertically concatenated into a combined matrix:

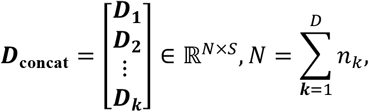

where *D* is the total number of recording days. In this concatenated matrix, each row corresponds to one neuron from one specific day, and each column corresponds to one regression feature (the same set of *S* features is used for all days).

#### 2. Joint SVD Decomposition

The normalized matrix is factorized via singular value decomposition:

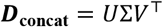

Where the matrix ***U*** represents coordinated patterns of neural activity within the population, while ***V*** encodes linear combinations of the eight stimulus conditions that are associated with specific neural response profiles.

#### 3. Dataset-Specific Projection

For each dataset ***D***_**K**_ in ***D***_**concat**_:

1. Let the index set *I_k_* identify the rows of U that belong to the neurons recorded on that day. The day-specific block of left singular vectors is:

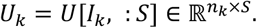
2. Perform QR decomposition to orthonormalize the basis:

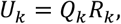

with sign alignment via correlation matching to ensure consistency. In QR decomposition, a matrix *U_k_* is factorized into the product of an orthogonal matrix *Q_k_* and an upper triangular matrix *R_k_*. *Q_k_* has orthonormal columns (i.e., *Q^T^Q* = *I*) and *R_k_* is upper triangular (all entries below the main diagonal are zero).
3. Project the centered data into the local subspace:

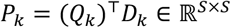

This framework facilitates the comparison of neural population dynamics across experiments, sessions, or individuals, while respecting the intrinsic variance structure of each dataset. It is particularly useful for identifying stable neural subspaces and tracking their reorganization over learning or time.

During conditioning phase, the framework is similar but the *S* = 6 (4 social demonstrator variable + 2 conditioning stimulus).

### Computation of Subspaces for Social Identities

To identify low-dimensional subspaces capturing the majority of social and location-related variance before and after conditioning, we utilized the regression coefficients, *β_i_*(*demo, loc*), obtained from joint SVD projection. Each social-location combination was represented at the population level by an 8-dimensional vector

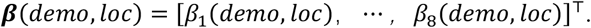

A demonstrator-specific subspace was defined as the set of vectors corresponding to the same demonstrator across different locations (ei. ***β***(*demo Reward, loc*1) and ***β***(*demo Reward, loc*2) form the demonstrator-Reward subspace). For each such group {*β*(*demo, loc*)}_*loc*=l,2_, we performed principal component analysis (PCA) to identify the first two orthogonal axes capturing the greatest response variance. This allowed the approximation:

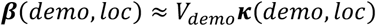

where 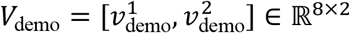 forms an orthogonal basis for the demonstrator-specific subspace, with 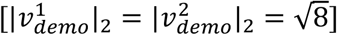and ***κ***(*demo, loc*) denotes the projection coefficients of ***β***(*demo, loc*) onto this subspace. For subsequent analysis, we used a normalized version of the basis, denoted 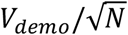 referred hereafter simply as 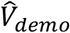 To facilitate comparison across different social subspaces, we used orthonormal axes 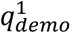and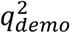rather than the original 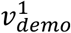 and 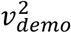to define each social subspace(*31*).

### Principal Angles Between Subspaces

To quantify the relative orientation of different social demonstrator subspaces in the neural state space during both pre- and post-conditioning phases, we computed the principal (or canonical) angles between subspaces—a geometric measure of alignment between two linear manifolds.

For any two demonstrator subspaces *a* and *b*, with corresponding orthonormal bases 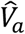 and 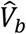, we performed singular value decomposition (SVD) on their inner product matrix:

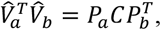

Where both *P_a_* and *P_b_* are 2×2 orthogonal matrices and *C* is a 2×2 diagonal matrix whose elements are the demo cosines of the principal angles *θ*_l_ and *θ*_2_:

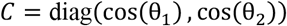

The principal angles θ_l_ ∈ [0, π/2] provide a measure of subspace similarity, with smaller angles indicating greater alignment(*31*).

During the conditioning phase, in which all demonstrators were presented at the same location within a single day, a simplified similarity measure was sufficient. Thus, we computed the cosine similarity and angle directly between subspace basis vectors without full principal angle analysis.

### The Variance Accounted For (VAF) Ratio

To further quantify the alignment between different social demonstrator subspaces, we introduced a variance-accounted-for (VAF) ratio. For a given demonstrator and location, let

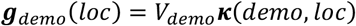

The VAF ratio between two demonstrator subspaces ***a*** and ***b*** is defined as:

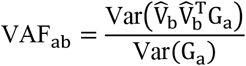

where

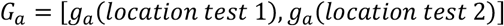

contains the reconstructed activity patterns for demonstrator a*a* across both test locations. Note that VAF_ab_ is asymmetric; it measures how well subspace ***b*** accounts for variance in the neural representation of demonstrator ***a***. The ratio ranges from 0 (orthogonal subspaces) to 1 (complete overlap)(*31*).

To compute the Variance Accounted For (VAF) between neural representations during the conditioning phase, we applied the following procedure:

#### 1. Data centering

Each dataset containing coefficients *β_i_*, obtained from L1+L2 linear regression about demonstrator variable and conditioning variable, was centered by subtracting its mean:

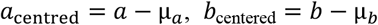

where *μ_α_* and *μ_b_* denote the means of datasets ***a*** and ***b***, respectively.

##### 2. Covariance Calculation

The covariance between the two centered datasets was computed as:

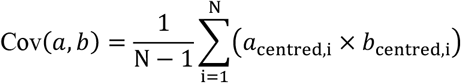

where ***N***=6, corresponding to the four demonstrators and two conditioning stimuli.

##### 3. Variance Estimation

The variances of the centered datasets were estimated using Bessel’s correction (ddof = 1) to account for sample bias:

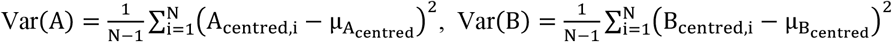

These quantities were subsequently used to compute the VAF ratio, quantifying the shared variance between neural activity patterns under different stimulus conditions.

##### 4. VAF computation

The Variance Accounted For (VAF) was then calculated as:

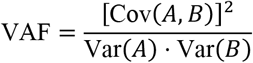

This metric quantifies the proportion of variance in one dataset that is explained by the other, yielding values between 0 (indicating no shared variance) and 1 (representing complete shared variance).

##### Gain-Modulation Approximation of Collective Variables

To assess whether a simplified gain-modulation mechanism could account for neural population responses, we approximated the full linear model at the level of collective variables using a gain-field modulation framework. Specifically, we modelled the latent variable ***κ***(*demo, loc*) as:

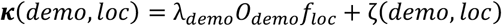

where *λ_demo_* is a demonstrator-specific gain factor, *f_loc_* (of length 2) is a spatial tuning vector independent of demonstrator identity, *O_demo_* (of size 2 ×2) is an orthogonal matrix aligning demonstrator and spatial features, and 𝜻(*demo, loc*) denotes the approximation error.

Following model fitting, gain factors were rescaled by the mean *λ* value across all conditions, and spatial vectors were adjusted accordingly to preserve relative modulation strength.

During the conditioning phase, where spatial variation was absent, the modulation index was defined as the Euclidean (L2) norm of the demonstrator-specific gain vector.

All modulation indices were z-scored across sessions prior to cross-day comparison to account for baseline shifts in response magnitude.

**Fig. S1.**
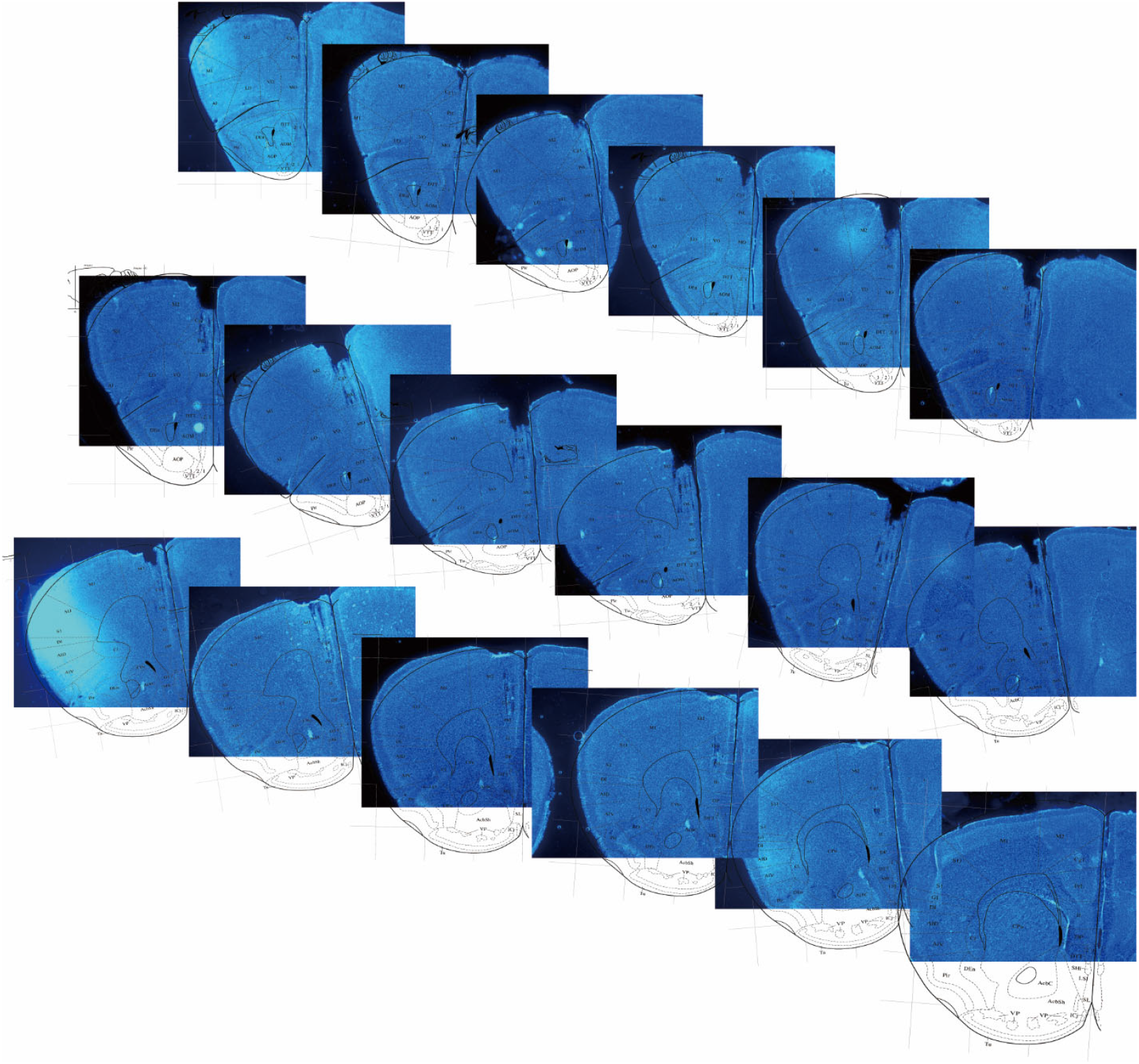
Histological verification of microelectrode tracks in mouse brain sections. Serial sections of mouse brain (spaced at 80 μm intervals). Image obtained from the Allen Mouse Brain Atlas (https://mouse.brain-map.org).

**Fig. S2.**
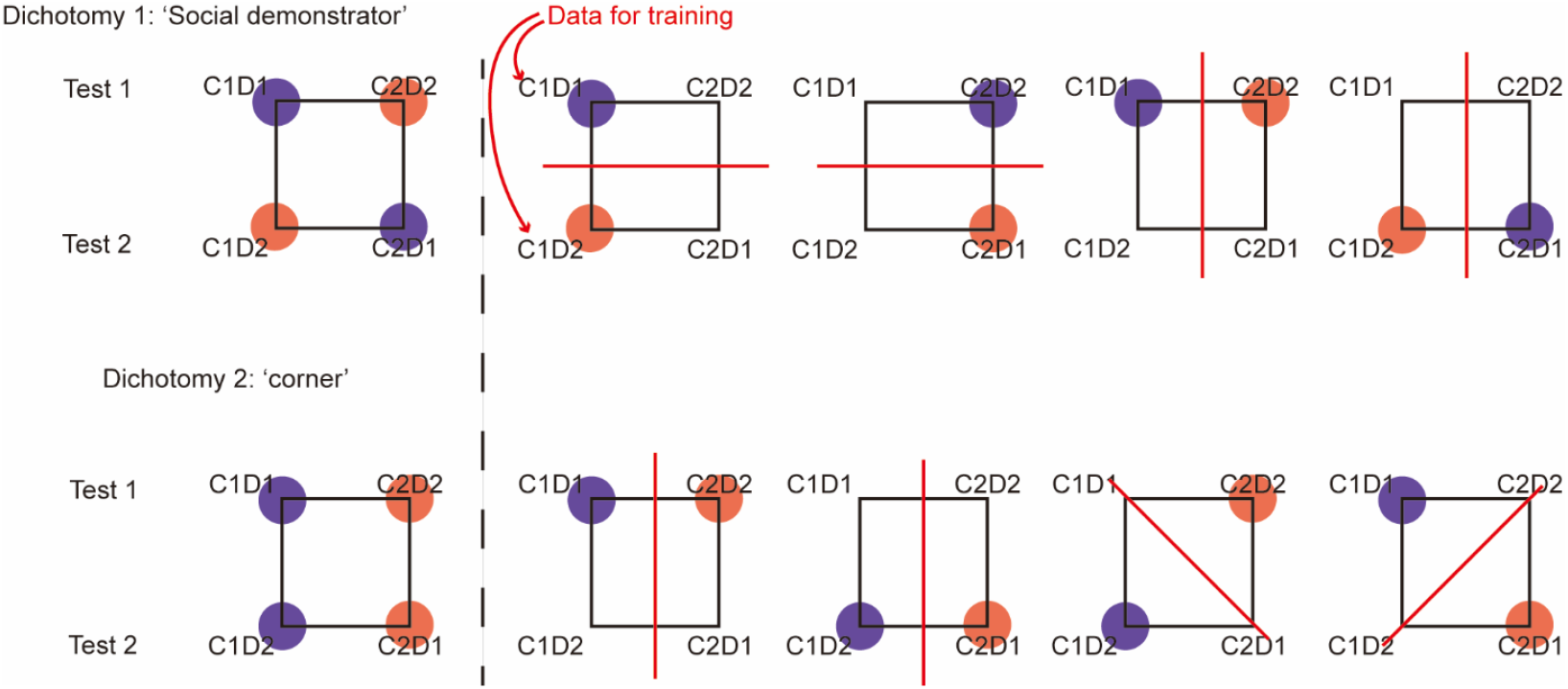
Explanation of the Computation Scheme for Cross-Condition Generalization Performance (CCGP). Each dichotomy corresponds to partitioning the four datasets into two groups (two datasets per group, represented by distinct colored clouds in the figure). All possible dichotomies correspond to well-defined variables (social demonstrator and positional corner). For each dichotomy, CCGP is computed by training a linear Support Vector Machine (SVM) on a subset of conditions (indicated by shaded colored circles in the figure) and subsequently testing it on the remaining conditions (non-shaded portion). For each dichotomy, there are four possible ways to select training conditions (choosing one condition from each side of the dichotomy), as illustrated in the row adjacent to each dichotomy in the figure. The final CCGP is obtained by computing the average test performance across all distinct training condition selection schemes. This schematic is simplified.

**Fig. S3.**
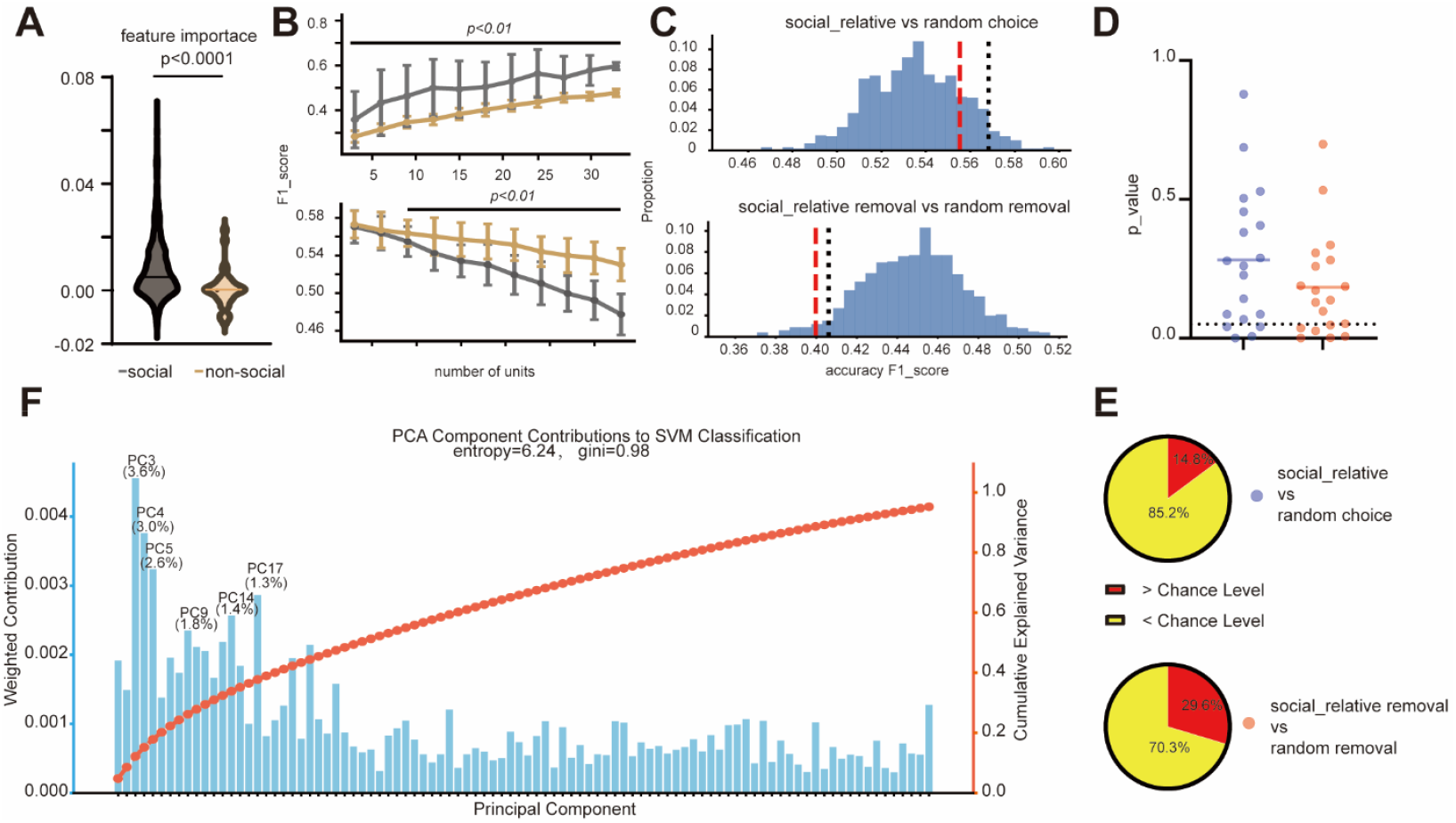
Social memory information encoded by most mPFC neurons is distributed in multiple neural orthogonal principal components. (**A**) Comparison of the feature importance values of these social-related neurons (including the pure social, conjunctive, and combinative neurons) *versus* those non-social neurons (including the corner and non-responsive neurons). (**B**) Changes of the classification accuracy in discriminating distinct demonstrators when the SVM model were trained with continuous additions (*top*) or ablations (*bottom*) of social-related neurons (yellow) and non-social neurons, respectively. *: p < 0.01, *t*-test. (**C**) Performance of control classifiers. F1 scores of support vector machine (SVM) classifiers trained on randomly selected (a) or randomly omitted (b) non-social neurons (with neuron numbers matched to social-related neurons; 1,000 iterations). The black dashed line indicates the 95% chance level threshold. The red line shows the true F1 score of social-related neurons. (**D**) Statistical significance of control tests. Distribution of p-values from the random selection test and the random omission test. (**E**) Proportion of above-chance classifications. The ratio of iterations yielding above-chance performance in the random selection (*top*) and random omission (*bottom*) tests. (**F**) Component analysis of population activity. Explained variance ratio and component weights after principal component analysis (PCA). The distribution of weights across many components, as indicated by Gini and entropy metrics, demonstrates that social discrimination is supported by a complex recognition process involving multiple, uncorrelated activity modes. For statistical details, see table S1.

**Fig. S4.**
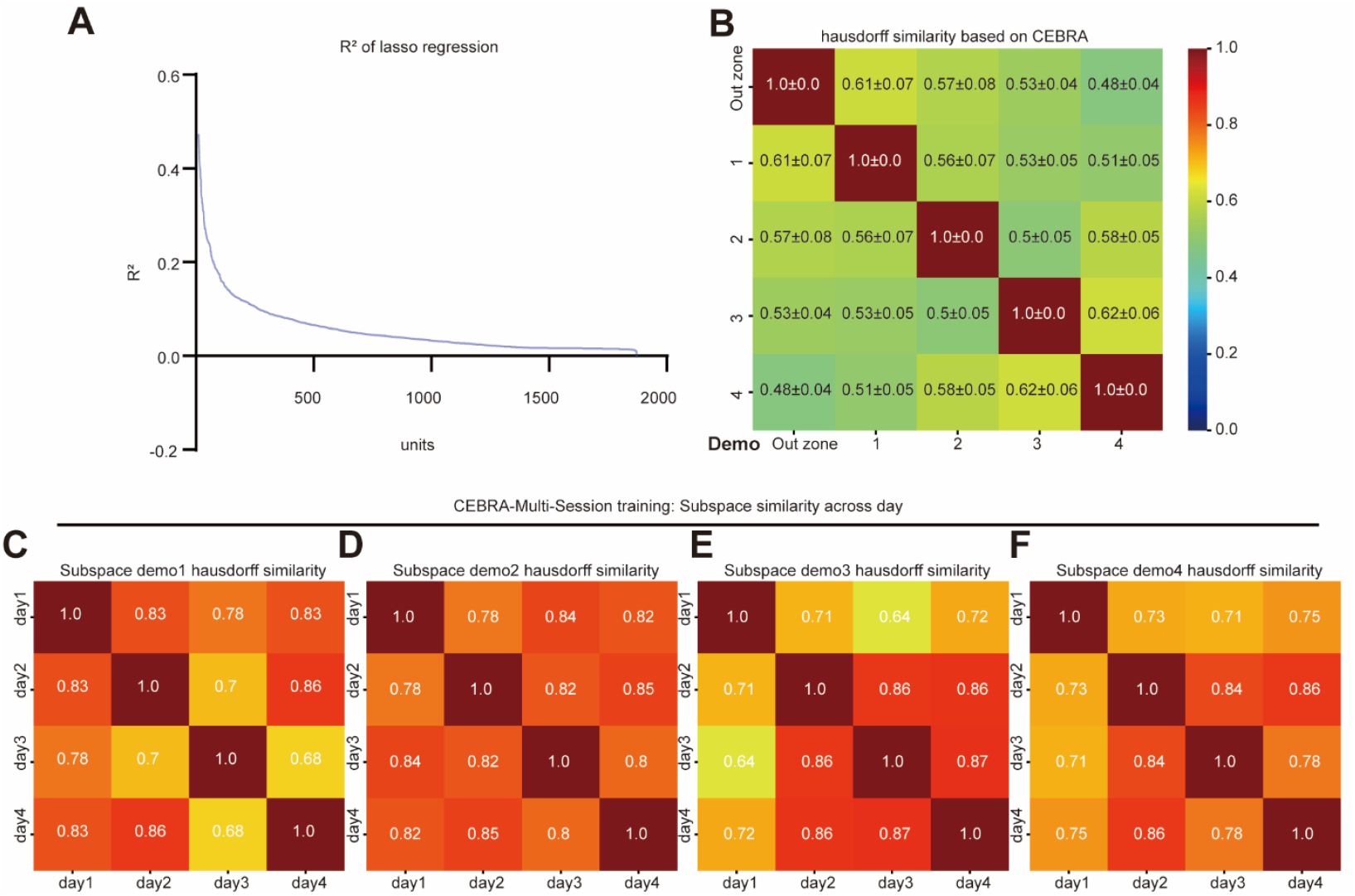
Stability of neural representations across days assessed via nonlinear dimensionality reduction and similarity analysis. (**A**) The ratio of variance in social discrimination explained by neural activity features selected via Lasso regression. (**B**) Similarity of neural representations across days, quantified using the Hausdorff similarity metric between neural embeddings generated by CEBRA (multi-session mode) for each pair of days. (**C** to **F**), Consistency of individual social subspaces. Similarity matrices (Hausdorff similarity) for each of the four identified social subspaces across the 4 recording days. For statistical details, see table S1.

**Fig. S5.**
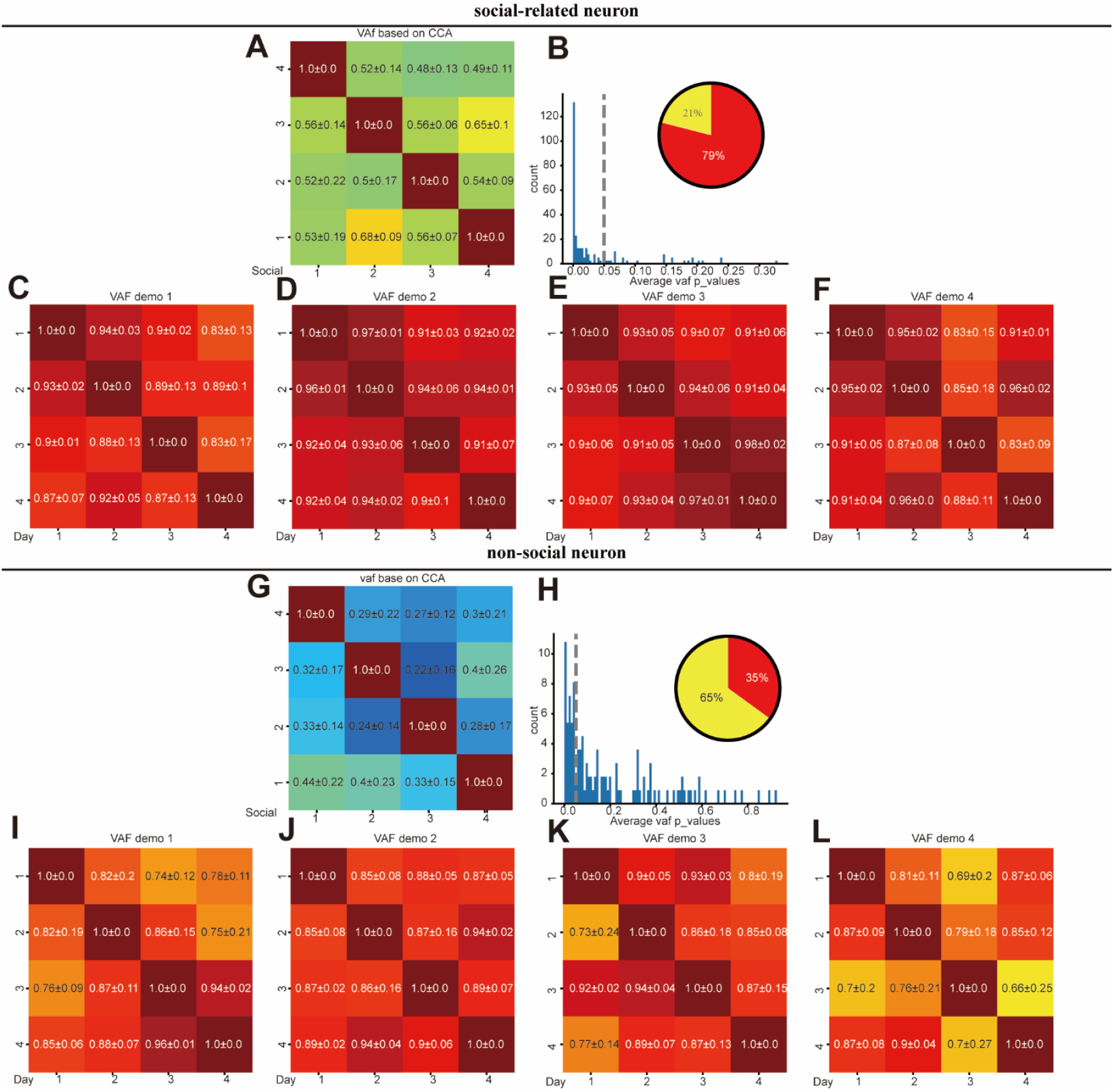
Social-related neurons exhibit higher representation stability cross days than non-social neurons. (**A**) VAF values were calculated between neural representations on different days using the dataset containing only social-related neurons. (**B**) Statistical significance of cross-day stability. Distribution of p-values from the comparison between the true VAF and VAF from shuffled data (1,000 iterations). Insets show the proportion of shuffled data points above (red) and below (yellow) the chance level. The dashed line indicates the significance threshold (α = 0.05). (**C** to **F**) Stability of individual social subspaces. VAF values for each of the four social subspaces across the 4-day period (using social-related neurons). (**G** to **L**) Similar analyses as (**A** to **F**), but using the dataset containing only non-social neurons. For statistical details, see table S1.

**Fig. S6.**
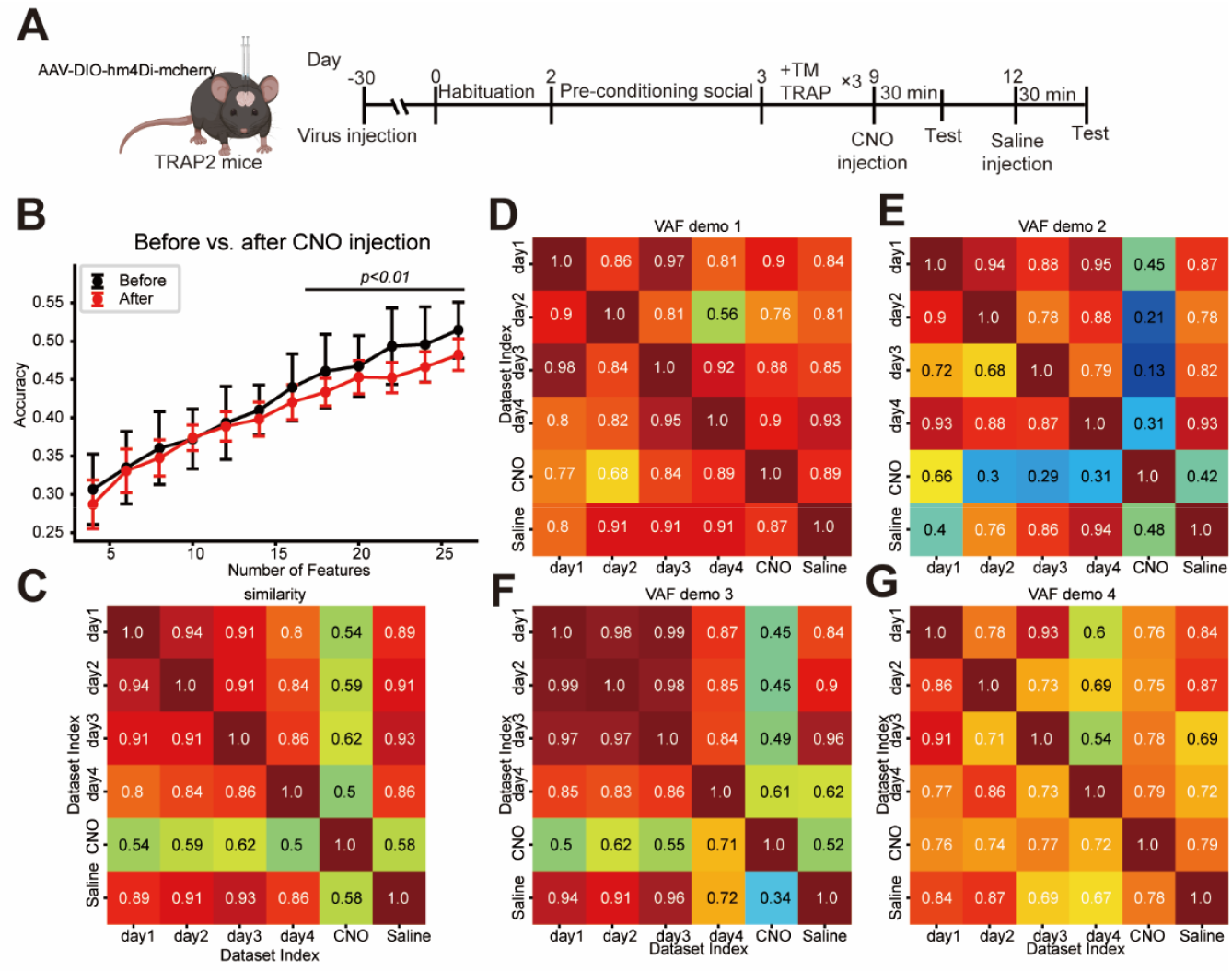
Chemogenetic suppression of behaviour-encoding neurons in the mPFC disrupts population representations. (**A**) Schematic of viral injection, cFos-TRAP labelling, and chemogenetic suppression (via DREADDs-hM4Di + Clozapine *N*-oxide, CNO) on neurons that are activated during the behavioural task. (**B**) Population decoding accuracy for social discrimination was significantly reduced after CNO injection to inhibit the TRAPed neuronal ensemble, compared to control conditions. (**C**) Neural activity from the chemogenetic suppression session failed to align within the same low-dimensional subspace as the embeddings from the untreated and control sessions, demonstrating a breakdown in representational structure. (**D** to **G**) Variance accounted for (VAF) ratio for each of the four social subspaces across days following chemogenetic suppression, showing disrupted stability. For statistical details, see table S1.

**Fig. S7.**
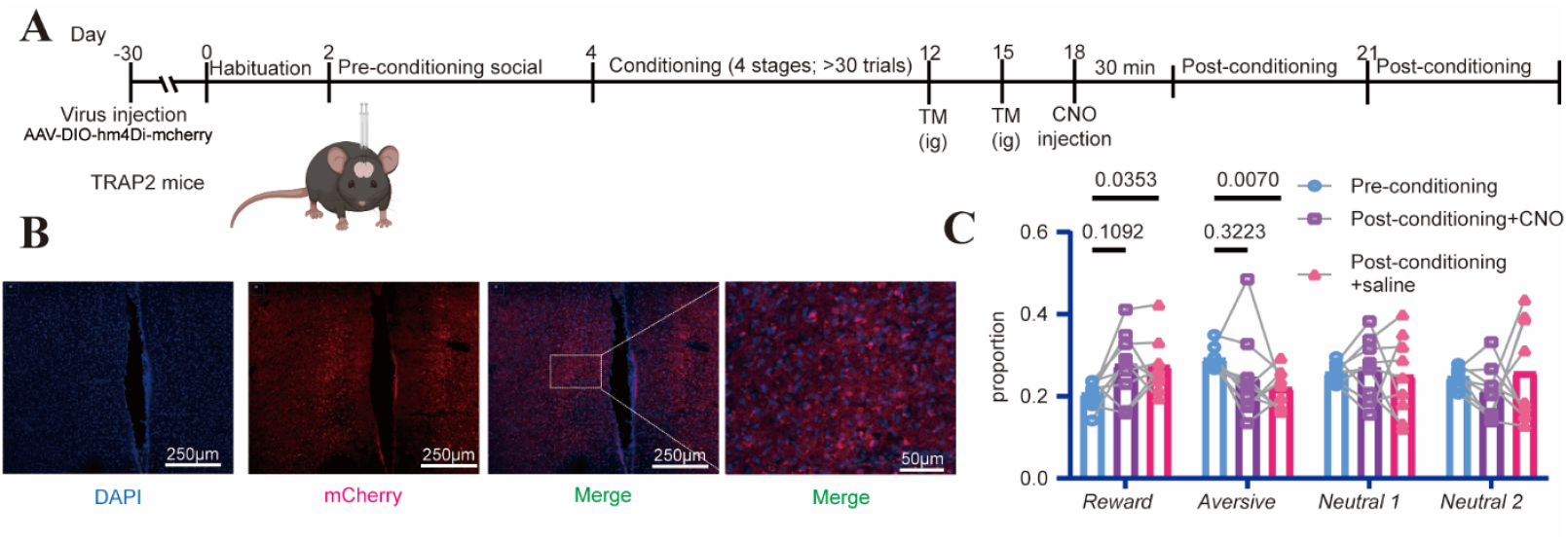
Chemo-genetic suppression of behaviour-encoding neurons in the mPFC impairs the social discrimination memory. (**A**) Schematic of the cFos-TRAP and chemogenetic suppression for targeting and manipulating neurons that are activated during the behavioral task. (**B**) A representative image showing the TRAPed neurons (expressing mCherry) in the mPFC. (**C**) Changes in the proportion of time spent investigating four individual social demonstrators across the following three different conditions: the Pre-conditioning, the Post-conditioning + CNO stage (social discrimination tests after the CNO injection) and the Post-conditioning + Saline (social discrimination tests after the saline injection). Chemogenetic suppression of the TRAPed neuronal ensemble significantly disrupted memory-dependent social investigations. For statistical details, see table S1.

**Fig. S8.**
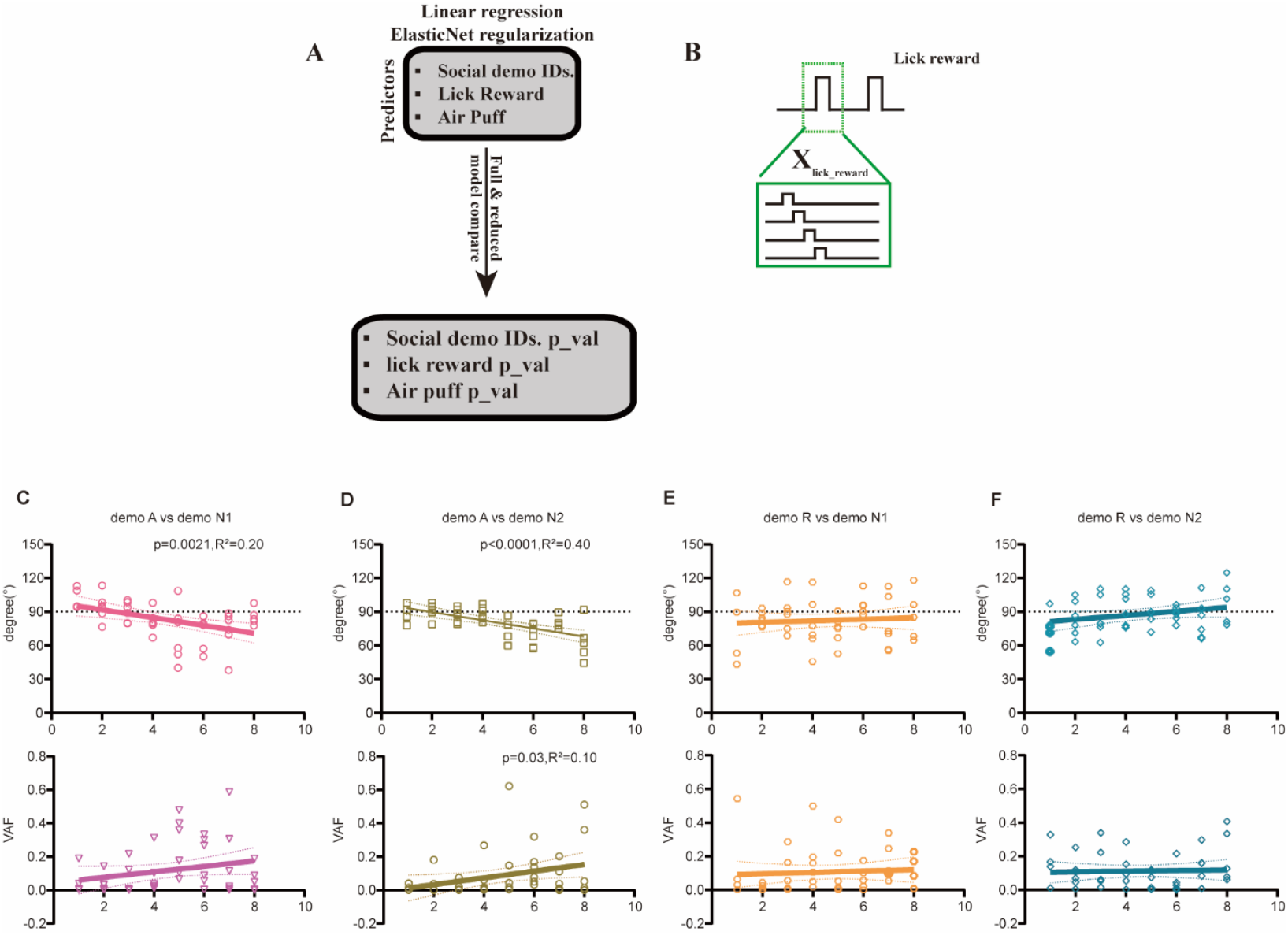
Dynamic reconfiguration of neural encoding toward the reward-paired demonstrator during conditioning. (**A**) Illustration of the linear regression framework with combined L1 (Lasso) and L2 (Ridge) regularization (Elastic Net) used to identify predictive neural features. (**B**) The temporally shifted design matrix was used to compute regression weights (kernels) for discrete behavioral events, isolating the neural components associated with the air puff (aversive, β_air puff_) and lick reward (milk reward, β_lick reward_). (**C** to **F**) Changes in the angles and variance accounted for (VAF) between selected subspaces during social conditioning. For statistical details, see table S1.

**Table S1.** Detailed statistical information.

| Figure | Sample size | Statistical test | Test statistic and P value |
| --- | --- | --- | --- |
| Fig. 1G | N=9 mice | Compare the MSE between full model and reduced model | threshold =0.01 |
| Fig. 1H | | Simple linear regression | *Non-response*: $R^2 = 0.009367$, $p\_value = 0.6242$;<br>*Corner neuron*: $R^2 = 0.04246$, $p\_value = 0.2928$;<br>*Social neuron*: $R^2 = 0.01613$, $p\_value = 0.5196$;<br>*Conjunctive neuron*: $R^2 = 0.03523$, $p\_value = 0.3388$;<br>*Combinative neuron*: $R^2 = 0.03219$, $p\_value = 0.3610$; |
| Fig. 2C | 9 mice<br>Number of pairs =26 | Paired t test | $t=9.526$, $df=25$<br>$p\_value < 0.0001$ |
| Fig. 2D | 9 mice | Simple linear regression | $F_{N1}=0.4965$, $p\_value_{N1}=0.4873$, $df_{N1}=1$, $n_{N1}=26$;<br>$F_{N2}=0.1931$, $p\_value_{N2}=0.6640$, $df_{N2}=1$, $n_{N2}=26$<br>$F_P=0.4417$, $p\_value_P=0.5121$, $df_P=1$, $n_P=26$;<br>$F_R=0.06502$, $p\_value_R=0.8007$, $df_R=1$, $n_R=26$;<br>$F_{all\_classify}=0.8273$, $p\_value_{all\_classify}=0.3714$, $df_{all\_classify}=1$, $n_{all\_classify}=26$; |
| Fig. 2E | 9 mice | CCGP | Shuffled CCGP=$0.5 \pm 0.02$ |
| Fig. 2F | 9 mice | SD | Shuffled SD=$0.5 \pm 0.02$ |
| Fig. 3G | 9 mice, show similar result. | Bootstrap shuffled | Generate the null-hypothesis distribution by shuffled one day of datasets across day, then compare the True value with this distribution. |
| Fig. 4B | N=20 | ANOVA | $F(2.330, 44.28) = 1.928$,<br>$P=0.1513$ |
| Fig. 4C | N=17 | Two-way ANOVA, Geisser-Greenhouse's correction Bonferroni's multiple comparisons test | *Reward* pre- vs. post, $t= 5.357$, Adjusted P Value = 0.0003, DF=16;<br>*Aversive* pre- vs. post, $t= 7.521$, Adjusted P Value <0.0001, DF=16;<br>*Neutral 1* pre- vs. post, $t= 1.235$, Adjusted P Value = 0.9384, DF=16;<br>*Neutral 2* pre- vs. post, $t= 1.044$, Adjusted P Value >0.9999, DF=16; |
| Fig. 4D | N=11 | Two-way ANOVA, Geisser-Greenhouse's correction Bonferroni's multiple comparisons test | *Reward* pre- vs. post, $t=4.317$, Adjusted P Value = 0.0061, DF=10;<br>*Aversive* pre- vs. post, $t=3.329$, Adjusted P Value = 0.0305, DF=10;<br>*Neutral 1* pre- vs. post, $t=0.05830$, Adjusted P Value >0.9999, DF=10;<br>*Neutral 2* pre- vs. post, $t=0.02518$, Adjusted P Value >0.9999, DF=10; |
| Fig. 4E | N=6 | Two-way ANOVA, Geisser-Greenhouse's correction Bonferroni's multiple comparisons test | *Reward* pre- vs. post, $t=0.6363$, Adjusted P Value = 0.9599, DF=5;<br>*Aversive* pre- vs. post, $t=1.356$, Adjusted P Value = 0.6541, DF=5;<br>*Neutral 1* pre- vs. post, $t=3.401$, Adjusted P Value = 0.0747, DF=5;<br>*Neutral 2* pre- vs. post, $t=0.2907$, Adjusted P Value = 0.9978, DF=5; |
| Fig. 4F | N=13 | Two-way ANOVA, Geisser-Greenhouse's correction Bonferroni's multiple comparisons test | *Reward* pre- vs. post, $t=3.932$, Adjusted P Value = 0.008, DF=12;<br>*Aversive* pre- vs. post, $t=3.520$, Adjusted P Value = 0.0169, DF=12;<br>*Neutral 1* pre- vs. post, $t=1.607$, Adjusted P Value = 0.5359, DF=12;<br>*Neutral 2* pre- vs. post, $t=1.650$, Adjusted P Value = 0.4995, DF=12; |
| Fig. 5A | 9 mice.<br>$N_{pre}=1866$,<br>$N_{post}=314$. | Chi-square test, Bonferroni's multiple comparisons test | $\chi^2(4)=10.30$, $p=0.036$;<br>*Non-responsive*, $Z=3.168$, Adjusted_p_value=0.0076;<br>*Corner* neuron, $Z=-0.803$, Adjusted_p_value=2.110;<br>*Social* neuron, $Z=0.028$, Adjusted_p_value=4.89;<br>*Conjunctive* neuron, $Z=-1.134$, Adjusted_p_value=1.28;<br>*Combinative* neuron, $stat=-1.671$, $p\_value=0.475$; |
| Fig. 5B | | | $\chi^2=12.12$, $p\_value=0.67$, $df=15$; |
| Fig. 5D to G | 6 mice.<br>$N_{pre}=1866$,<br>$N_{post}=314$. | Generate distribution by random subsample same amount from pre-conditioning dataset to SVM, 1000 times. | $p_{Reward}=0.028$;<br>$p_{Aversive}=0.039$;<br>$p_{Neutral\ 1}=0.177$;<br>$p_{Neutral\ 2}=0.086$; |
| Fig. 5I | N=6 mice. | ANOVA, Tukey's multiple comparisons test | *Reward* vs. *Aversive*, pre- vs. post-conditioning, $p=0.04$, $q=3.029$, DF=30.<br>*Reward* vs. *Neutral 1*, pre- vs. post-conditioning, $p=0.30$, $q=1.490$, DF=30. |
|  |  |  | *Reward* vs. *Neutral 2*, pre- vs. post-conditioning, $p=0.31$, $q=1.463$, $DF=30$.<br>*Aversive* vs. *Neutral 1*, pre- vs. post-conditioning, $p=0.33$, $q=1.400$, $DF=30$.<br>*Aversive* vs. *Neutral 2*, pre- vs. post-conditioning, $p=0.26$, $q=1.613$, $DF=30$.<br>*Neutral 1* vs. *Neutral 2*, pre- vs. post-conditioning, $p=0.61$, $q=0.72$, $DF=30$. |
| Fig. 5J | N=6 mice. | ANOVA, Tukey's multiple comparisons test | *Reward* vs. *Aversive*, pre- vs. post-conditioning, $p=0.03$, $q=3.279$, $DF=30$.<br>*Reward* vs. *Neutral 1*, pre- vs. post-conditioning, $p=0.10$, $q=2.396$, $DF=30$.<br>*Reward* vs. *Neutral 2*, pre- vs. post-conditioning, $p=0.10$, $q=2.389$, $DF=30$.<br>*Aversive* vs. *Neutral 1*, pre- vs. post-conditioning, $p=0.13$, $q=2.203$, $DF=30$.<br>*Aversive* vs. *Neutral 2*, pre- vs. post-conditioning, $p=0.05$, $q=2.936$, $DF=30$.<br>*Neutral 1* vs. *Neutral 2*, pre- vs. post-conditioning, $p=0.16$, $q=2.032$, $DF=30$. |
| Fig. 5K | N=6 mice. | ANOVA, Tukey's multiple comparisons test | *Reward*, pre- vs. post-, $p=0.0084$, $q=5.950$, $df=5$;<br>*Aversive*, pre- vs. post-, $p=0.58$, $q=0.8373$, $df=5$;<br>*Neutral 1*, pre- vs. post-, $p=0.24$, $q=1.90$, $df=5$;<br>*Neutral 2*, pre- vs. post-, $p=0.72$, $q=0.54$, $df=5$; |
| Fig. 5M | N=6 mice. | ANOVA, Tukey's multiple comparisons test | *Reward* vs. *Aversive*, pre- vs. post-, $p=0.009$, $q=3.807$, $DF=66$;<br>*Aversive* vs. *Reward*, pre- vs. post-, $p=0.0133$, $q=3.598$, $DF=66$;<br>*Reward* vs. *Neutral 1*, pre- vs. post-, $p=0.9154$, $q=0.1507$, $DF=66$; |
|  |  |  | *Neutral 1* vs. *Reward*, pre- vs. post-,<br>$p=0.2071$, $q=1.802$, $DF=66$;<br>*Reward* vs. *Neutral 2*, pre- vs. post-,<br>$p=0.3407$, $q=1.357$, $DF=66$;<br>*Neutral 2* vs. *Reward*, pre- vs. post-,<br>$p=0.0809$, $q=2.507$, $DF=66$;<br>*Aversive* vs. *Neutral 1*, pre- vs. post-,<br>$p=0.009$, $q=3.793$, $DF=66$;<br>*Neutral 1* vs. *Aversive*, pre- vs. post-,<br>$p=0.014$, $q=3.573$, $DF=66$;<br>*Aversive* vs. *Neutral 2*, pre- vs. post-,<br>$p=0.014$, $q=3.569$, $DF=66$;<br>*Neutral 2* vs. *Aversive*, pre- vs. post-,<br>$p=0.0109$, $q=3.706$, $DF=66$;<br>*Neutral 1* vs. *Neutral 2*, pre- vs. post-,<br>$p=0.1075$, $q=2.308$, $DF=66$;<br>*Neutral 2* vs. *Neutral 1*, pre- vs. post-,<br>$p=0.0453$, $q=2.885$, $DF=66$; |
| Fig. 6A to D |  | T test between in zone and out zone. | Validated by Linear regression. |
| Fig. 6E to F | N=7 | Simple linear regression | *Social* response: $R^2=0.03261$,<br>$p=0.3674$;<br>*Condition* response: $R^2=0.1279$, $p=0.067$;<br>*Joint* response: $R^2=0.3628$,<br>$p\_value=0.0009$;<br>Demo *Reward* response: $R^2=0.018$,<br>$p=0.4647$;<br>Demo *Aversive* response: $R^2=0.00014$,<br>$p=0.8403$;<br>Demo *Neutral 1* response: $R^2=0.018$, $p=0.4647$;<br>Demo *Neutral 2* response: $R^2=0.021$, $p=0.4277$; |
| Fig. 6 H to J | N=6 | Simple linear regression | Milk reward:<br>$R^2=0.1915$, $p\_value=0.0026$;<br>Air puff: |
| | | | $R^2=0.3967$, $p\_value<0.0001$;<br>*Demo Reward*:<br>$R^2=0.2249$, $p\_value=0.001$;<br>*Demo Aversive*:<br>$R^2=0.02693$, $p\_value=0.2814$;<br>*Demo Neutral 1*:<br>$R^2=0.041$, $p\_value=0.1829$;<br>*Demo Neutral 2*:<br>$R^2=0.0089$, $p\_value=0.5371$; |
| Fig. 6K<br>to L<br>fig. S8 C<br>to F | N=6 | Simple linear<br>regression | *Angle demo Reward* vs. *demo Aversive*:<br>$R^2= 3.178e-005$, $p= 0.9707$;<br>*Angle demo Reward* vs. *demo Neutral 1*:<br>$R^2= 0.006382$, $p= 0.6019$;<br>*Angle demo Reward* vs. *demo Neutral 2*:<br>$R^2= 0.06305$, $p= 0.1002$;<br>*Angle demo Reward* vs. *milk reward*:<br>$R^2= 0.02230$, $p= 0.3276$;<br>*Angle demo Aversive* vs. *Neutral 1*:<br>$R^2= 0.1994$, $p= 0.0021$;<br>*Angle demo Aversive* vs. *Neutral 2*:<br>$R^2= 0.3985$, $p < 0.0001$;<br>*Angle demo Aversive* vs. *air puff*:<br>$R^2= 0.003333$, $p = 0.7064$;<br>*Angle demo Neutral 1* vs. *demo Neutral 2*:<br>$R^2= 0.0002$, $p = 0.9266$;<br>*Angle Milk Reward* vs. *Air puff*:<br>$R^2= 0.04698$, $p = 0.1527$;<br>*VAF demo Reward* vs. *demo Aversive*:<br>$R^2= 9.424e-005$, $p= 0.9495$;<br>*VAF demo Reward* vs. *demo Neutral 1*:<br>$R^2= 0.004401$, $p= 0.6650$;<br>*VAF demo Reward* vs. *demo Neutral 2*:<br>$R^2= 0.001495$, $p= 0.8032$;<br>*VAF demo Reward* vs. *milk reward*:<br>$R^2= 3.820e-005$, $p= 0.9679$;<br>*VAF demo Aversive* vs. *Neutral 1*:<br>$R^2= 0.06084$, $p= 0.1024$;<br>*VAF demo Aversive* vs. *Neutral 2*:<br>$R^2= 0.1047$, $p = 0.0321$;<br>*VAF demo Aversive* vs. *air puff*: |
| | | | $R^2 = 0.1799, p = 0.0037$;<br>VAF demo *Neutral 1* vs. demo *Neutral 2*:<br>$R^2 = 0.002736, p = 0.7360$;<br>VAF Milk *Reward* vs. Air puff:<br>$R^2 = 0.009260, p = 0.5295$; |
| fig. S2A | 9 mice.<br>N<sub>social</sub>=1120 ,<br>N<sub>non-social</sub>=624 | Mann Whitney test | $p < 0.0001$,<br>Mann-Whitney U=272516 |
| fig. S2B | | T test | Random subsample or remove same number of neurons, 50 times,<br>$p$ value threshold=0.01 |
| fig. S2C to E | 9 mice | Random removal or random involve |  |
| fig. S2F |  | PCA to 95% explain ratio, then calculate SVM weight, gini and entropy. | High entropy means the progress recruit many principal dimensions;<br>High gini means weight value is contributed from few principal dimensions;<br>This result means the social discrimination is complex progress, because it recruit many dimension but main on few dimension |
| fig. S3A | 9 mice | $R^2$ after linear regression | |
| fig. S3B to F | 9 mice | Hausdorff similarity (Method) |  |
| fig. S4 | 9 mice, show similar result. Separated social relative neuron and non-social relative neurons. | Bootstrap shuffled | Generate the null-hypothesis distribution by shuffled one day of datasets across day, then compare the True value with this distribution. |
| fig. S5 | | T test | Random subsample or remove same number of neurons, 50 times,<br>$p$ value threshold=0.01 |
| fig. S6 | N=7mice | Two-way ANOVA, Geisser-Greenhouse's correction Bonferroni's multiple comparisons test | Demo *Reward*:<br>Pre-cond. VS. CNO:<br>Adjusted $p = 0.1170$, $q = 3.221$, $DF = 8$;<br>Pre-cond. VS. Saline:<br>Adjusted $p = 0.0218$, $q = 4.850$, $DF = 8$;<br>Demo *Aversive*: |
|  |  |  | Pre-cond. VS. CNO:<br>Adjusted $p$= 0.6932, $q$= 1.181, DF=8;<br>Pre-cond. VS. Saline:<br>Adjusted $p$= 0.0129, $q$= 5.372, DF=8;<br>*Demo Neutral 1:*<br>Pre-cond. VS. CNO:<br>Adjusted $p$= 0.8748, $q$= 0.7027, DF=8;<br>Pre-cond. VS. Saline:<br>Adjusted $p$= 0.9360, $q$= 0.4919, DF=8;<br>*Demo Neutral 2:*<br>Pre-cond. VS. CNO:<br>Adjusted $p$= 0.0889, $q$= 3.487, DF=8;<br>Pre-cond. VS. Saline:<br>Adjusted $p$= 0.9870, $q$= 0.2180, DF=8; |
| fig. S7C | | Chi-square test | $\chi^2(1) = 395.8, p < 0.0001$ |

**Movie S1. Pre-conditioning test 1 video**

Type or paste caption here.

**Movie S2. Pre-conditioning test 2 video**

Type or paste caption here.

**Movie S3. Reward Conditioning video**

Type or paste caption here.

**Movie S3. Aversive Conditioning video**

Type or paste caption here.

**Movie S5. Post-conditioning test 1 video**

Type or paste caption here.

**Movie S6. Post-conditioning test 2 video**

Type or paste caption here.

